# A centimeter-long bacterium with DNA compartmentalized in membrane-bound organelles

**DOI:** 10.1101/2022.02.16.480423

**Authors:** Jean-Marie Volland, Silvina Gonzalez-Rizzo, Olivier Gros, Tomáš Tyml, Natalia Ivanova, Frederik Schulz, Danielle Goudeau, Nathalie H Elisabeth, Nandita Nath, Daniel Udwary, Rex R Malmstrom, Chantal Guidi-Rontani, Susanne Bolte-Kluge, Karen M Davies, Maïtena R Jean, Jean-Louis Mansot, Nigel J Mouncey, Esther Angert, Tanja Woyke, Shailesh V Date

## Abstract

Cells of most bacterial species are around 2 µm in length, with some of the largest specimens reaching 750 µm. Using fluorescence, x-ray, and electron microscopy in conjunction with genome sequencing, we characterized *Ca.* Thiomargarita magnifica, a bacterium with an average cell length greater than 9,000 µm that is visible to the naked eye. We found that these cells grow orders of magnitude over theoretical limits for bacterial cell size through unique biology, display unprecedented polyploidy of more than half a million copies of a very large genome, and undergo a dimorphic life cycle with asymmetric segregation of chromosomes in daughter cells. These features, along with compartmentalization of genomic material and protein synthesis in membrane-bound organelles, indicate gain of complexity in the *Thiomargarita* lineage, and challenge traditional concepts of bacterial cells.

**One Sentence Summary:** *Ca*. T. magnifica are compartmentalized centimeter-long bacteria

## Main Text

Bacteria and archaea are the most diverse and abundant organisms on Earth. With only a small fraction of them isolated in culture, we remain grossly ignorant of their biology (*1*). While most model bacteria and archaea are small, some remarkably large cells, referred to as giant bacteria, are evident in at least four phyla (*2*), and have cellular sizes in the range of tens or even hundreds of microns (*3, 4*). Some exceptional members of sulfur-oxidizing gammaproteobacteria *Thiomargarita namibiensis,* for instance, are known to reach up to 750 µm (average size: 180 µm) (*4–6*). Such bacterial giants raise the question of whether more macro-bacteria might still be out there but have not yet been identified.

Here, we describe a novel sessile filamentous *Thiomargarita* species from a marine sulfidic environment that dwarfs all other known giant bacteria by about 50-fold. Our multi-faceted imaging analyses reveal massive polyploidy and a dimorphic developmental cycle where genome copies are asymmetrically segregated into apparent dispersive daughter cells. Importantly, we show that centimeter-long *Thiomargarita* filaments represent individual cells with genetic material and ribosomes compartmentalized into a novel type of membrane-bound organelle. Sequencing and analysis of genomes from five single cells revealed insights into distinct cell division and cell elongation mechanisms. These unique cellular features likely allow the organism to grow to an unusually large size and circumvent some of the biophysical and bioenergetic limitations on growth. In reference to its exceptional size, we propose to name this species *Thiomargarita magnifica* (referred to below as *Ca.* Thiomargarita magnifica).

### Ca. Thiomargarita magnifica is a centimeter-long, single bacterial cell

Some Large Sulfur Bacteria (LSB) form very long filaments which may reach several centimeters in length, but they are composed of thousands of individual cells which do not exceed 200 µm (*7–10*). Here we observed seasonal “bouquets” of centimeter-long white filamentous *Thiomargarita* cells attached to sunken leaves of *Rhizophora mangle* (Fig. S1) in shallow tropical marine mangroves from Guadeloupe, Lesser Antilles. *Thiomargarita* spp. are sulfur-oxidizing gammaproteobacteria known to be morphologically diverse and display striking polyphenism (*11*). The morphology of the filaments observed in Guadeloupe resembled those of sessile *Thiomargarita*-like cells reported from deep-sea methane seeps (*12*). They had a stalk-like shape for most of their length and constricted gradually towards the apical end forming buds (Figs. 1A-E). In contrast to relatives that live buried in sediment, these filaments were smooth in appearance and free of epibiotic bacteria or any extracellular mucus matrix (Fig. S2) (*11*). Budding filaments had an average length of 9.72 ± 4.25 mm, and only the most apical constrictions closed completely to form 1-4 rod-shaped separate cells of 0.21 ± 0.05 mm. We also noted some filaments reaching a length of 20.00 mm (Figs. 1A, S1, S3), much larger than any previously described single-celled prokaryote.

**Fig. 1.**
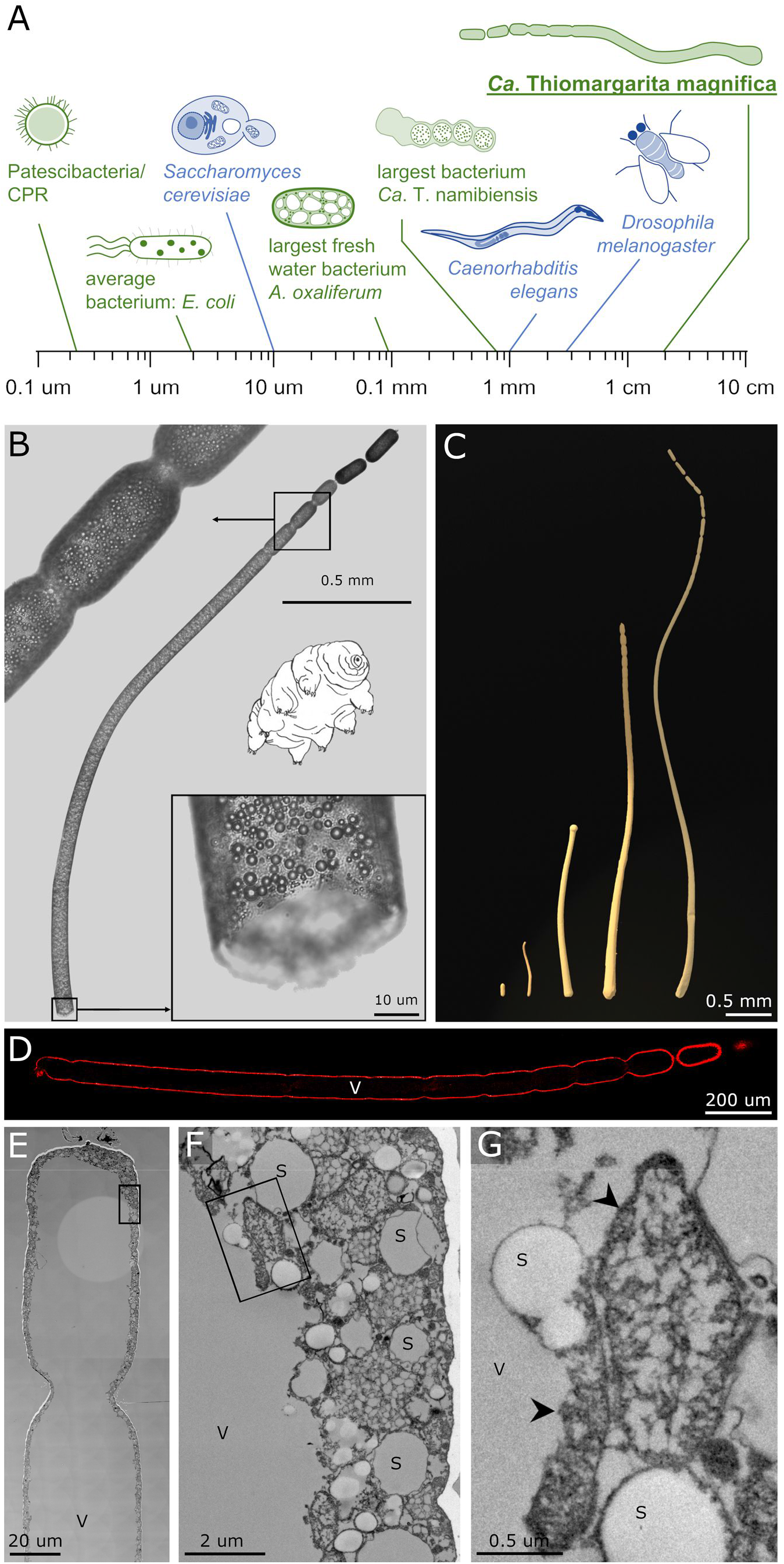
Morphology and ultrastructure of *Ca.* Thiomargarita magnifica. A: Size comparison of selected bacterial (green) and eukaryotic (blue) model systems on a log scale. B: Light microscopy montage of the upper half of a *Ca.* T. magnifica cell, with a broken basal part revealing a tube-like morphology due to the large central vacuole and numerous spherical intracellular sulfur granules (a tardigrade is shown for scale). C: 3D rendering of segmented cells from HXT (Movies S1, S2 and S6) and CLSM (Movie S3), putatively at various stages of the developmental cycle. From left to right 3D rendered cells are cell *D*, *B*, *F*, *G*, and *D* (Table S2). Note smallest stage correspond to cell *D* terminal segment and was added to the left for visualization purposes. D: CLSM observation of cell *K* after fluorescent labeling of membranes with FM 1-43x showing the continuity of the cell from the basal pole to the first complete constriction at the apical end. E: TEM montage of the apical constriction of a cell, with the cytoplasm constrained to the periphery. F: Higher magnification of the area marked in E, with sulfur granules and pepins at various stages of development. G: Higher magnification of the area marked in E showing two pepins (arrowheads). S: sulfur granule; V: vacuole.

To further characterize *Ca.* T. magnifica cells, we highlighted membranes using osmium tetroxide or fluorescent dye FM 1-43x and visualized entire filaments in 3D with Hard X-ray Tomography (HXT, n=4) and Confocal Laser Scanning Microscopy (CLSM, n=6) as well as filament sections (up to 850.6 µm long) with Transmission Electron Microscopy (TEM, n=15). Strikingly, all techniques consistently showed that each filament was one continuous cell for nearly its entire length with no division septa, including the partial constrictions towards the apical pole. Only the most apical few buds were separated from the filament by a closed constriction and represented daughter cells (Figs. 1, S5 and Movies S1-S4).

Similar to other Large Sulfur Bacteria (LSB), *Ca.* T. magnifica cells showed a large central vacuole which was continuous along the whole filament and accounted for 73.2 ± 7.5 % (n=4) of total volume (Figs. 1D, E, S5 and Table S2). The cytoplasm was 3.34 ± 1.48 µm thick and was constrained to the periphery of the cell (Figs. 1E, F and S5). TEM revealed numerous electron lucent vesicles 2.40 ± 1.03 µm in diameter which corresponded to the refractile granules observed with bright-field microscopy and represented sulfur granules as shown by Energy Dispersive X-ray Spectroscopy (Figs. 1F, S5, S10 and Supplementary Text). The cell envelope consisted of a thick outer layer covering the cytoplasmic membrane (Fig. S5). The cytoplasm of *Ca*. T. magnifica appeared to be compartmentalized in the form of dense regions similar to other LSB (*7, 10, 13*), comprising multiple membrane-bound bodies 1.31 ± 0.70 µm in diameter (Figs. 1F, G). We hypothesized that some of these dense regions within the cytoplasm, which were distinct from large sulfur granules, may contain dispersed genomic material, as polyploidy is evident in many giant bacteria (*2, 14*).

### Ca. T. magnifica DNA is contained in a novel type of membrane-bound bacterial organelle

While bacteria were once presumed to be un-compartmentalized “bags of enzymes,” recent studies show the presence of organelles with functions as diverse as anaerobic ammonium oxidation, photosynthesis or magnetic orientation (*15*). No bacteria or archaea are known to unambiguously segregate their genetic material in the manner of eukaryotes, although some evidence for a possible membrane-bound DNA compartment occupying most of the cell’s volume in a member of the Atribacteria has been reported (*15, 16*). Surprisingly, DAPI staining revealed DNA in *Ca*. T. magnifica cells was concentrated in membrane-bound granules (Fig. 2H-K), and not spread throughout the cytoplasm, as is common in bacteria. These DNA containing bodies also harbored electron-dense structures of 10 to 20 nm in size, similar to the signature of ribosomes (Fig. 2F-G). Fluorescence *in situ* hybridization (FISH) with probes specifically targeting ribosomal RNA sequences of *Thiomargarita* confirmed that ribosomes were indeed present and concentrated in these membrane-bound structures (Figs. 2 and S6 to S8), which were spread throughout the entire cell, including the apical buds (Fig. S8). This compartmentalization of DNA and ribosomes is reminiscent of genomic compartmentalization in eukaryotes and represents a novel cellular structure within bacteria. By analogy with pips – the numerous small seeds in fruits such as watermelon or kiwi – we propose to name this bacterial organelle a *“pepin”* (singular *pepin*, plural *pepins*: from vulgar latin *pép*, an expressive creation used to express smallness).

**Fig. 2.**
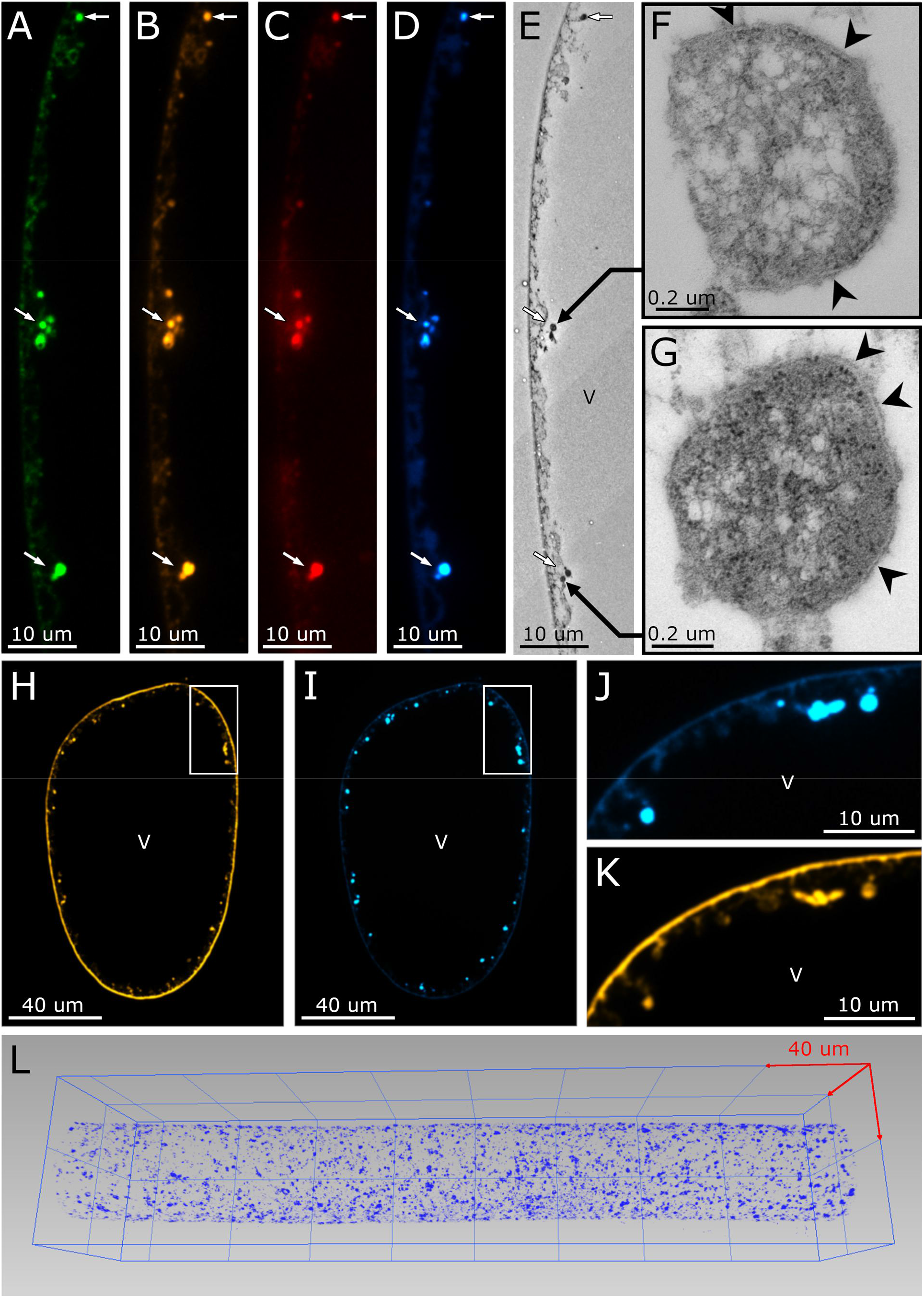
Characterization of the pepin organelles by Fluorescent In Situ Hybridization (FISH) and correlative Transmission Electron Microscopy (TEM) as well as membrane and DNA staining. A-D: Pepins (arrows) in the cytoplasm of *Ca*. T. magnifica (class gammaproteobacteria) are labeled with the general bacterial probe EUB labeled with Alexa Fluor® 488 (A, green), gammaproteobacteria specific probe Gam42a labeled with Cy3 (B, yellow), *Thiomargarita-*specific probe Thm482 labeled with Cy5 (C, red) and with DAPI (D, blue) (see Supplementary Text for details). E: TEM of a serial thin section consecutive to the semi-thin section used for FISH. The FISH and DAPI positive pepins appear as electron dense organelles under TEM. F and G: Pepins (from E) under higher magnification; pepins are delimited by a membrane (arrowheads) and contain numerous ribosomes which appear as small electron dense granules. H and I: Fluorescent labeling of membranes using FM 1-43x (H) and DNA using DAPI (I) on a cross section of a cell. The pepins labeled with DAPI are also labeled with the dye FM 1-43x confirming the presence of a membrane. J and K: Higher magnifications of the area delimited by the white rectangle in H and I. L: 3D visualization of a central portion of a cell after DAPI staining (blue) showing the multitude of DNA clusters spread throughout the cytoplasm (cell *M,* Table S2 and Movie S5).

### A highly polyploid cell with a large genome

All previously described giant bacteria are polyploid (*2, 3, 14*), *i.e.* their cells contain large numbers of genome copies – ranging from tens to tens of thousands – dispersed throughout the cell, supporting the local need for molecular machineries and overall cellular growth (*17, 18*). Polyploidy has been shown to decrease the selective pressure on genes, allowing intracellular gene duplication, reassortment and divergence and lead to extreme intracellular genetic diversity in some LSB (*19*). On the other hand, it may also allow balancing of genome copies through homologous recombination and support a high level of genome conservation (*20*). *Ca*. T. magnifica like all bacterial giants appeared to be polyploid; counts of DAPI stained DNA clusters on three CLSM 3D dataset suggested an average of 36,880 ± 7,956 genome copies per millimeter of filament (737,598 ± 159,115 for a fully grown 2 cm cell, see Table S2, Fig. 2L, Movie S5 and details in Supplementary Text). This is the highest estimated number of genome copies for a single cell. It is one order of magnitude above that of other giant bacteria (*2, 18*).

To genomically characterize *Ca.* T. magnifica, we amplified, sequenced, and assembled the DNA of five individual cells collected from a single sunken leaf (Tables S3-S4). All five draft genomes appeared highly similar to each other with an Average Nucleotide Identity (ANI) above 99.5% (Table S5). Variant analysis within the single-cell genome sequences indicated a genomically homogenous population (1.22 ± 0.18 SNPs/100 kbp, Table S6) (*21*), which is similar to other polyploid bacteria (*20, 22*). The assemblies were estimated to be near complete at 91.0% to 93.7% with total sequence lengths between 11.5 Mb and 12.2 Mb. This is twice as large as the only other sequenced *Thiomargarita* species *Ca.* T. nelsonii (*23, 24*) and at the upper range of bacterial genome sizes; bacterial genomes are on average 4.21 ± 1.77 Mb (Fig. S9). The *Ca*. T. magnifica genome from filament 4 contained 11,788 genes (only half with a functional annotation, see Table S4), more than three times the median gene count of prokaryotes (3,935 genes) (*25*). For the sake of comparison with eukaryotic organisms, *Ca*. T. magnifica has a genome as large as the baker yeast *S. cerevisiae* (12.1 Mb) and contains more genes than the model fungus *Aspergillus nidulans* (≍ 9,500 genes).

Analysis of the genome revealed a large set of genes for sulfur oxidation and carbon fixation, suggesting chemoautotrophy, in accordance with our other evidence for thioautotrophy (Fig. S10, S11 and Supplementary Text). Like its sister lineage *Ca.* T. nelsonii, *Ca*. T. magnifica encoded a wide range of metabolic capabilities with one remarkable difference: it lacked nearly all genes involved in dissimilatory and assimilatory nitrate reduction, and denitrification except for Nar and Nap nitrate reductases. This suggests that nitrate can solely be used as an electron acceptor (*23, 24*) (see Supplementary Text for extended genome analysis). The somewhat surprising absence of epibiotic bacteria, despite its size, may be explained by the high number of genes encoding secondary metabolism. With 25.9% of sequences dedicated to biosynthetic gene clusters (Fig. 3A), the genome encoded dozens of modular Non-Ribosomal Peptide Synthetases (NRPSs) and Polyketide Synthases (PKSs) systems, hinting at numerous secondary metabolism pathways (similar to Actinobacteria (Table S7)), that are indicative of antibiotic or bioactive compound production.

**Fig. 3.**
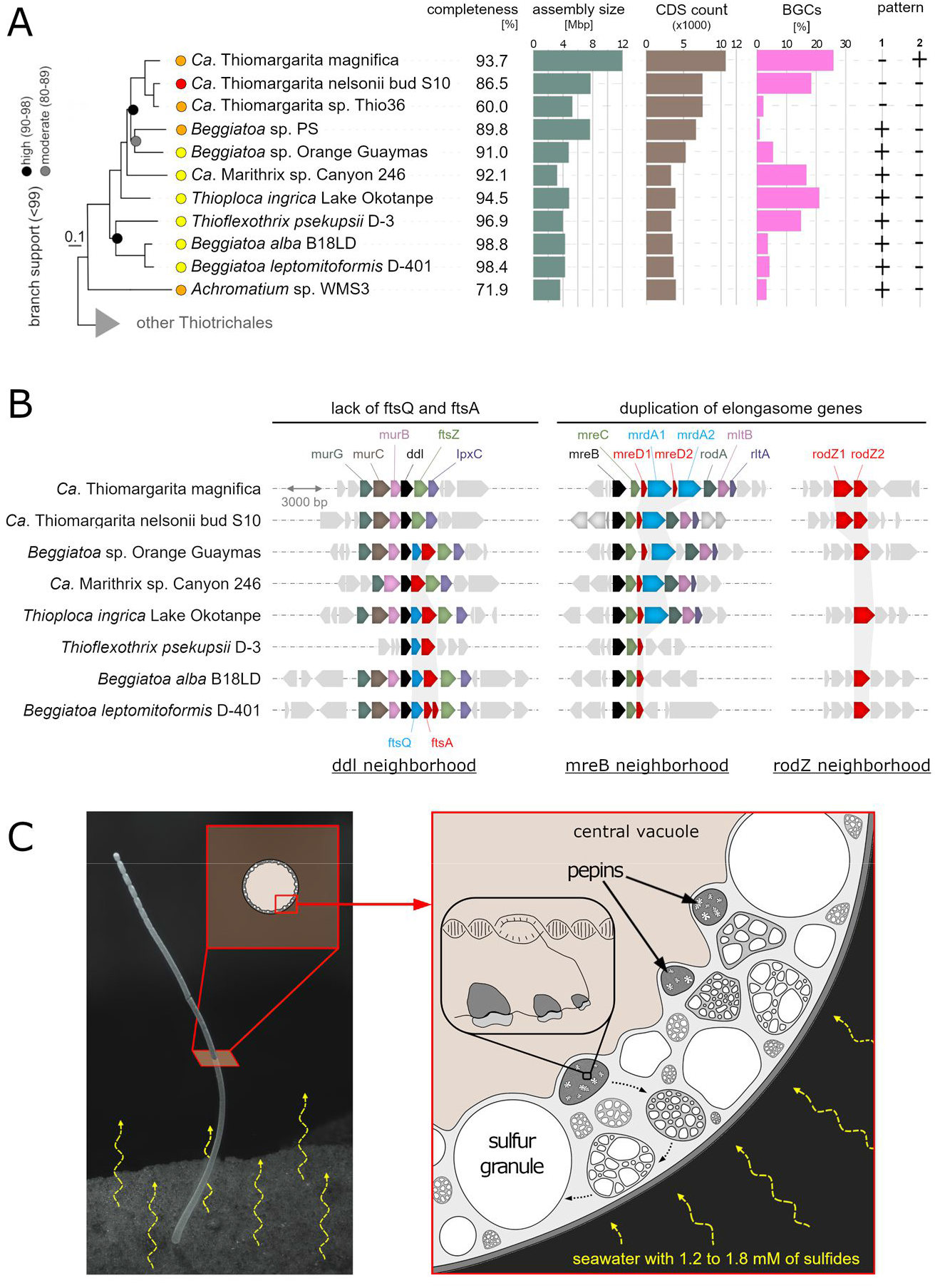
Genome analysis and proposed model for the sub-cellular organization of *Ca.* Thiomargarita magnifica. A. Genome phylogenetic tree with added information about genome quality (red: low quality, orange: medium quality, and yellow: high quality (*40*)), estimated level of completeness, assembly size, CDS count, and percentage of sequence dedicated to biosynthetic gene clusters (BGCs). Pattern 1 corresponds to “complete gene cluster for cell division of model bacteria”. Pattern 2 corresponds to “*mreD*, *mrdA* and *rodZ* genes are duplicated”. B. Gene neighborhoods centered on the *ddl*, *mreB* and *rodZ* genes showing the incomplete set of divisome genes (lack of *ftsQ* and *ftsA*) in both *Thiomargarita* species as well as the duplication of elongasome genes (*mreD*, *mrdA* and *rodZ*) in *Ca.* T. magnifica. Note that *Beggiatoa* sp. PS, *Achromatium* sp. WMS3 and *Ca*. Thiomargarita sp. Thio36 draft genomes were too fragmented and were not included here. C. Light microscopy image and model proposed for the sub-cellular organization in *Ca.* Thiomargarita magnifica showing how the pepin organelles greatly increase the surface area of putative bioenergetic membranes.

The *Ca.* T. magnifica genome also held clues for its unusual cell morphology, with an atypical complement of cell division and cell elongation genes. Many genes that encode core cell division proteins, including core components of Z ring assembly and regulation, FtsA, ZipA and FtsE-FtsX, were lacking (Fig. 3B, Table S8), whereas genes that encode the cytoskeletal protein FtsZ, which is part of well-conserved dcw (“division and cell wall”) operon and the core component of Z ring, and proteins ZapA, ZapB and ZapD, which interact with FtsZ and regulate Z ring assembly, were conserved (*26*). Even more remarkably, the entire set of genes that encode late divisome proteins, including peptidoglycan polymerases FtsI and FtsW, as well as FtsQ, FtsL, FtsB, and FtsK, was absent from all *Ca.* Thiomargarita genomes (Fig. 3B, S16 and Table S8). This conspicuous lack of cell division genes was contrasted by a complete set of genes encoding cell elongation proteins, three of which - *mreD*, *rodZ* and peptidoglycan transpeptidase *mrdA*- have undergone recent duplications, with both copies located next to each other on the chromosome (Fig. 3B, S13-15, Table S8) (*26*). It is possible that an increased number of cell elongation genes, coupled with the lack of key cell division genes, may be responsible for producing the unusually long filaments of *Ca.* T. magnifica (see Supplementary Text).

### Dimorphic developmental cycle of Ca. T. magnifica

Laboratory observations of live *Ca.* T. magnifica revealed eventual apical bud detachment from the filament and release into the environment, likely representing a dispersive stage of the developmental cycle (Fig. 1C; S1B-F and Supplementary Text). We observed dozens of cells at all intermediary stages from the smallest attached cells resembling terminal segments recently settled to the largest filaments with apical constrictions (Fig. S1 and Movies S1-S3 and S6). Such a dimorphic life cycle resembles the aquatic single-celled model system *Caulobacter crescentus* as well as the multicellular segmented filamentous bacteria, albeit at a different scale, in which stalked “parent” cells produce free living “daughter” cells (*27, 28*). Due to this asymmetrical division mode, only a small fraction of the genome copies – present within pepins in the most apical bud - were transmitted to the daughter cell (Fig. S8). Like the polyploid giant bacterium *Epulopiscium* spp., *Ca.* T. magnifica apparently transmits only a subset of its genomes, to be called ‘germ line genomes,’ to the offspring (*14, 18*). If terminal buds are indeed daughter cells, such a developmental cycle may have evolved to enhance dispersion similar to the fruiting bodies of the social myxobacteria or to the aerial hyphae of *Streptomyces* spp. (*29*). This apparent life cycle is also somewhat analogous to the sulfur oxidizing giant ciliate symbiosis, *Zoothamnium niveum* (*30*), possibly representing a case of convergent evolution of developmental cycle across domains (see Supplementary Text).

### Concluding remarks

While cells of most bacteria and archaea are around 2 µm, eukaryotic cells are usually between 10 and 20 µm with some of the largest single-cell eukaryotes reaching 3 to 4 cm (*31*). Several theoretic frameworks explain the restriction of bacteria and archaea to microscopic sizes: i) lack of active intracellular transport and the reliance on chemical diffusion, which is efficient only along micrometer distances (*4*); ii) a predicted maximum cell volume constraining the number of needed ribosomes should the cell grow larger (*32*); or even iii) a decrease in energy efficiency due to mismatched surface area to volume ratio when considering placement of membrane-bound ATP synthases (*33*). These frameworks all suggest that with increasing size, the metabolic needs of a bacterial cell grow faster than the cell’s capacity to sustain it and should reach a limit. The next largest prokaryote known after *Ca.* T. magnifica, - *Ca.* T. nelsonii - has a metabolically active biovolume of 1.05×10^-14^ m^3^ (excluding the central vacuole), close to the predicted maximum due to ribosome limitations: 1.39xyn10^-15^ m^3^ (*32*). Our precise 3D measurements on a 4.27 mm *Ca.* T. magnifica cell revealed a cytoplasm biovolume three orders of magnitude above that limit (5.91×10^-12^ m^3^, Table S2). It is possible, as may be the case with *Ca.* T. magnifica, that changes in spatial organization of cellular components and rearrangement of the bioenergetic membrane system may allow some bacteria to overcome many limitations (Fig. 3C).

The origin of biological complexity is among the most important, yet unanswered, questions in biology. While most bacteria are considered small and simple, some have evolved complex innovations. Functionally diverse bacterial microcompartments are found in at least 23 phyla (*34*). Cyanobacteria can form multicellular centimeter long filaments and are capable of cell differentiation (*35*). Planctomycetes have special energy transduction organelles called anammoxosomes, a compartmentalized cell and some are even capable of phagocytosis (*15, 36*). The social Myxobacteria have large genomes, a complex developmental cycle and are capable of moving and feeding cooperatively in predatory groups (*37*). Through its gigantic cell size, its large genome, its di-morphic life cycle, but most importantly through its compartmentalization of genetic material in membrane-bound pepins, *Ca.* T. magnifica adds to the list of bacteria that have evolved a higher level of complexity. It is the first and only bacteria known to date to unambiguously segregate their genetic material in membrane-bound organelles in the manner of eukaryotes and therefore challenges our concept of a bacterial cell.

Confirmation bias related to viral size prevented the discovery of giant viruses for more than a century, and their ubiquity is only now being recognized (*38, 39*). The discovery of *Ca.* T. magnifica suggests that large and more complex bacteria may be hiding in plain sight. Investigating their rare biology, energy metabolism and the precise role and nature of pepins, will take us a step closer in understanding the evolution of biological complexity.

## Supplementary Materials

Materials and Methods

Supplementary text

Figures S1-S16

Tables S1-S11

Movies S1-S6

References (*1–77*)

## Acknowledgments

We are thankful for the following centers where the electron microscopy analyses were performed: i) the *Centre Commun de Caractérisation des Matériaux des Antilles et de la Guyane* in Guadeloupe, F.W.I., ii) the *Electron Microscopy Lab* (EML) of the University of California Berkeley, iii) the *Electron Microscopy Resource in Donner* at LBNL, Berkeley, and iiii) the *FEI Eindhoven center*. We are particularly grateful to Daniel Jorgens at EML for advice and assistance in electron microscopy sample preparation and data collection. The x-ray tomography data were acquired at the *Stanford Nano Shared Facilities* in Stanford University, Palo Alto, California and we are particularly grateful to Arturas Vailionis for his technical support during HXT scan acquisitions. The confocal microscopy observations were performed at the *Advanced Microscopy Facility* at LBNL, Berkeley, California. Preliminary confocal microscopy observations were acquired at the *IBPS imaging facility* which is supported by “Conseil Regional d’Ile-de-France”. We are grateful to Sebastien Volland for his help with 3D rendering animations and to Heather Maughan for English editing of this manuscript.

## Author contributions following the CRediT model

Conceptualization: OG, SGR, JMV, TW, SVD, FS, RRM, NI, TT, NHE

Methodology: OG, TT, JMV, SVD, SGR, DG, NN, RRM, TW, CGR, SBK

Formal analysis: JMV, SGR, FS, NI, DU, OG

Investigation: JMV, SGR, OG, TT, FS, DU, DG, NN, NI, CGR, SBK, NHE, MRJ

Data curation: NI, FS, JMV, DU, SGR

Visualization: OG, SGR, JMV, FS, NI, DU

Resources: KMD, TW, RRM, OG, JLM, NHE, NJM

Supervision: OG, TW, SVD

Funding acquisition: TW, SVD, OG, JLM

Writing – original draft: JMV

Writing – review & editing: JMV, SGR, OG, TW, SVD, EA, NI, RRM

## Supplementary Materials

Materials and Methods

Supplementary text

Figures S1-S16

Tables S1-S11

Movies S1-S6

References (*1–77*)

## Supplementary Materials for

**This PDF file includes:**

Materials and Methods

Supplementary Text

Figs. S1 to S16

Tables S1 to S11

Captions for Movies S1 to S6

**Other Supplementary Materials for this manuscript include the following:**

Movies S1 to S6

### Materials and Methods

#### Sampling

Samples of Large Sulfur Bacteria (LSB) were collected from a marine mangrove environment (ambient temperature 28°C) in “La manche à eau” in Guadeloupe, Lesser Antilles, at one site (16°16’40”N, 61°33’28”W) (*41*). Sunken leaves of *Rhizophora mangle* containing LSB were sampled by hand from the surface layer of the sediment (*c.* 1 m depth). Living LSB samples were processed within 1 h after collection under a dissecting microscope. The samples were then washed with 0.22 μm filtered seawater prior to use for molecular experiments. Individual bacteria were fixed at 4°C either for 4 hours in 4% paraformaldehyde in sterile seawater, or in 2.5% glutaraldehyde in 0.1 M cacodylate buffer (pH 7.2), which was made iso-osmotic (900 mOsmoles) with sea water by the addition of sodium chloride and calcium chloride. Samples were stored at 4°C until analysis.

#### Light microscopy

Samples were observed live or fixed under standard stereomicroscopes. If applicable, a series of images captured at different focus distances was merged using the focus stacking software Helicon Focus® (Fig. S1). Fixed samples were also observed using a light microscope Axio Observer.D1 (Carl Zeiss, Jena, Germany) equipped with a black-and-white high-resolution camera (AxioCam MRm, Carl Zeiss, Jena, Germany).

#### Ultrastructural analysis

For conventional SEM analysis, samples were briefly rinsed in the cacodylate buffer and then dehydrated through a graded acetone series before drying under CO2 using a critical point dryer machine (EM CPD300, Leica). The samples were then sputter-coated with gold (Sputter Coater SC500, BioRad) before observation at 20 kV with an FEI Quanta 250 scanning electron microscope.

For scanning transmission electron microscopy (STEM) analysis, prefixed bacterial filaments were washed twice in 0.1 M sodium cacodylate buffer to remove aldehydes before fixation in 1% osmium tetroxide for 45 min at room temperature. Samples were then rinsed in distilled water and post-fixed with 2% aqueous uranyl acetate for 1 h at room temperature. After three washes with distilled water, each sample was dehydrated through a graded acetone series and embedded in Epon-Araldite according to Glauert (1975) (*42*). Thin sections (60 nm thick) were stained for contrast for 30 min in 2% aqueous uranyl acetate before examination with a Quanta 250 (FEI - STEM Mode).

For Transmission Electron Microscopy (TEM) observations, glutaraldehyde fixed samples were washed in a cacodylate buffer and post-fixed with osmium tetroxide as described above. They were then cryo-immobilized using a BAL-TEC HPM 010 high-pressure freezer and further placed in a freeze-substitution medium made of 1% osmium tetroxide, 0.1% uranyl acetate, and 5% ddH2O in acetone. The samples were freeze-substituted following the super quick procedure described in McDonald (2011) (*43*). The substitution medium was washed away with pure acetone and the samples were infiltrated and flat-embedded in Epon-Araldite resin as described in Müller-Reichert et al. (2003) (*44*). Thin sections of 70 nm were mounted on formvar coated slot grids and stained for 4 min in 2 % uranyl acetate followed by 4 min in Reynolds Lead Citrate. Slot grids were observed with a FEI Tecnai 12 or a Jeol 1400FLASH TEM. Montages were acquired either manually or automatically using the SerialEM software. Tile images were assembled either manually using GIMP, or with the Etomo program from the Imod suite (*45*), or with the software Image Composite Editor (Microsoft), or with the FIJI package on ImageJ (*46*). Morphometric measurements were performed using FIJI. The average thickness of the cytoplasm was obtained from 118 measurements realized on sections from three different cells (cell 1: n = 51; cell 2: n = 17; cell 3: n = 50). The average diameter of the elemental sulfur granules was obtained from 338 measurements realized on sections from three different cells (cell 1: n = 103; cell 2: n = 31; cell 3: n = 204). The average diameter of pepins was obtained from 92 measurements realized on sections from five different cells (cell 1: n = 8; cell 2: n = 29; cell 3: n = 20; cell 4: n = 5; cell 5: n = 30).

#### Elemental analysis

The elemental composition of fully hydrated samples was analyzed using an Environmental Scanning Electron Microscope (ESEM). The dehydration step was avoided because the elemental sulfur S8 is soluble in alcohol and acetone. The samples were fixed in 2.5% glutaraldehyde in seawater and kept in the same solution until examination. Samples were simply washed quickly in distilled water to remove salts and then introduced to an ESEM (FEI Quanta FEG) under a pressure of 650 Pa. ESEM studies were carried out using various acceleration voltages to reveal the presence of sulfur in the samples, as well as their morphology. We used 1) 15 kV for back scattered electron images (Z contrast) and energy dispersive X-ray spectroscopy (analyses and elemental mapping) and 2) 3 kV for secondary electron images (sample morphology).

#### Confocal Laser Scanning Microscopy (CLSM)

Paraformaldehyde-fixed *Ca.* T. magnifica cells (n=8) were washed three times in sterile seawater, dehydrated through an ascending ethanol series - to dissolve elemental sulfur granules - and rehydrated through a descending ethanol series. Individual cells were transferred onto glass slides equipped with Gene Frame (Thermofisher) to avoid crushing the cells with the coverslips. Membrane labeling of cells was carried out using the lipophilic dye FM 1-43x (Molecular Probes, Eugene, OR, USA) according to the manufacturer’s protocol. DNA was labelled with DAPI (4′,6-Diamidino-2-phenylindole dihydrochloride; Millipore Sigma) following the manufacturer’s protocol. We imaged 8 cells on a Zeiss LSM 710 microscope by acquiring multiple overlapping Z-stack images (tiles) with the Zen software (Zeiss). From these 8 cells, 6 were imaged in their entire length (cells F to K, see Table S2) and were used to confirm the single cell nature of the filaments, measure precise morphometric parameters (length, minimum and maximum diameters), and assess polyploidy level for one of them. The remaining 2 cells were imaged only partially (cells L and M, see Table S2) and used exclusively to assess the polyploidy level by counting DAPI signal clusters. As positive controls, we followed the same protocol and prepared multicellular filaments of the Cyanobacteria *Microcoleus vaginatus* from a pure culture, *Beggiatoa-*like filaments collected in a Guadeloupe mangrove, and *Marithrix*-like filaments collected at the White Point Beach hydrothermal vent (*47, 48*) (Fig. S4; Movie S4). All 3D data sets were imported into the ORS Dragonfly software for stitching, 3D rendering and morphometric analyses and/or polyploidy analysis. Three cells observed in their entirety with high lateral resolution (cells F, G and H) were segmented using the ORS Dragonfly deep learning tool and manual segmentation tools. Precise volumes of the central vacuole and the cytoplasm were measured (see Table S2).

#### Hard x-ray computed tomography

We used hard x-ray computed tomography to visualize six cells in three dimensions with isotropic resolution. Four cells were observed in their entirety and two were observed only partially (see Table S2). *Ca.* T. magnifica cells fixed with glutaraldehyde and post-fixed with osmium tetroxide, as described above, were washed three times in sterile seawater, dehydrated through ascending ethanol series and stored in 70 % ethanol at 4° C. Before analysis, cells were re-hydrated and immobilized inside plastic capillaries with 1 % low melting point agarose. After sealing the capillaries on both ends, we glued them onto the head of a sewing pin and imaged them in a Zeiss Xradia 520 Versa x-rays microscope. The technical details of each scan are provided in Table S1. The CT scans were reconstructed using the Zeiss software and further imported in the ORS Dragonfly software for stitching of the tiles, segmentation and analysis. Four cells were observed in their entirety (cells A, B, D and E), three of which were segmented using the ORS Dragonfly deep learning tool and manual segmentation tools. Precise total volumes of the cells were measured and for one of them (cell D) the volumes of the cytoplasm and central vacuole were measured as well (see Table S2).

#### Fluorescent in situ hybridization (FISH) and TEM correlation

Paraformaldehyde fixed cells (n=4) were washed in seawater before dehydration through an ascending ethanol series. We then infiltrated the cells with medium grade LR-White resin. LR-White embedded cells were polymerized at 40 °C under anaerobic conditions for three days. We analyzed with FISH a total of 84 semi-thin sections (500 nm), coming from 7 different areas analyzed in triplicates within each of the four cells. Sections were mounted on PTFE coated microscope glass slides (Electron Microscopy Sciences, Hatfield, PA) and analyzed with FISH. Some consecutive thin sections (70 nm) were prepared for correlative TEM as described in the ultrastructural analysis section. The FISH hybridization solution (0.9 mol L-1 NaCl, 20 mmol L-1 Tris/HCl pH 8.0, 0.01% SDS, 10% formamide) was applied onto the section in a 20 µL drop containing 0.5 µM of each oligonucleotide probe. Hybridization was performed in a humid chamber at 46°C for 3 h. Washing was performed under stringent conditions at 48°C for 15 min (*49*). We used a combination of a eubacterial probe mixture (EUB338 Alexa Fluor® 488 single labeled) (*50*), a general gammaproteobacteria probe (Gam42a Cy3 double-labeled) (*49*), an unlabeled competitor probe (BET42a) (*49*) and a genus-specific *Thiomargarita* probe (Thm482, Cy5 double-labeled) (*51*). Nonsense probes (Non-EUB) (*52*) labeled in Cy3 and Cy5 were also applied to all slides to control for false positive signals due to autofluorescence or nonspecific probe binding, but no signals were observed in these controls. We mounted the FISH slides with antifadent solution CitiFluor AF1 Plus DAPI (Electron Microscopy Sciences, Hatfield, PA). Micrographs were taken using a 63x oil-immersion objective on an Inverted epifluorescence microscope (Axio Observer.D1, Carl Zeiss, Jena, Germany) equipped with a black-and-white high-resolution camera (AxioCam MRm, Carl Zeiss, Jena, Germany). FISH images were overlaid with their corresponding TEM observations in the GIMP software.

#### Sulfide measurements

Sunken leaves with attached elongated cells were brought to the laboratory and placed in a mesocosm mounted the day before measurement to simulate their mangrove environment. The samples were collected from the field on the day of measurement. Sulfide measurements were carried out using H2S100 microsensors (Unisense®) attached to a micromanipulator (type MD4 Rechts, Märzhäuser®). One microsensor was placed into the water column 5 cm above the bacterial cells and the other positioned in the middle of the LSB “bouquet” attached onto a submerged leaf of *Rhizophora mangle*. The measurements were recorded every 30 s using SensorBasic® software. Calibrations were performed according to the Unisense® instructions. The pH was measured with an autonomous probe (NKE), similar to that described by Le Bris *et al.*, (2001) (*53*), fixed to the micromanipulator. Total sulfide concentrations (S^2-^tot= H2S+HS^-^+S^2-^) were calculated, accounting for the measured pH and salinity using a pK of 6.51 (*54*).

#### Genome sequencing, assembly, binning and annotation

We processed five *Ca.* T. magnifica filaments for single-cell genomic sequencing. Within one hour after sampling, we dissected individual cells out of the decaying leaf and washed them three times in sterile seawater before storing them at −80°C. We thawed each individual filament and immediately amplified the genomic DNA by multiple displacement amplification using the REPLI-g kit (Qiagen). DNA libraries were created from 200pg of DNA from each of the amplified products using Nextera XT DNA library creation kit (Illumina). We sequenced the DNA libraries on an Illumina Nextseq High Output platform. We then imported pair-end reads (2×150 bp) into the KBase platform (www.kbase.us) (*55*). In KBase, we used SPAdes (v3.13.0) to assemble reads into contigs of at least 500 bp (using kmers of 33, 67, 99, 125 bp). We then binned contigs over 2000 bp using MetaBAT2 resulting in 2 to 6 bins per filament (see Supplementary Table 3). Only one bin per filament was taxonomically identified as *Thiomargarita* by the GTDB-Tk classify app (v0.1.4). We extracted the contigs from the *Thiomargarita* bin and treated them as an assembly for further analyses (referred to as: filament n *Ca*. T. magnifica genome). We assessed genome qualities with CheckM (v1.0.18).

#### Genome analysis

##### Average Nucleotide Identities and clonality

We choose filament #5 as a reference genome based on its better assembly statistics. We computed pairwise Average Nucleotide Identities (ANIs) with FastANI. We assessed within filament clonality by mapping the reads from filament #1, #2, #3 and #4 onto filament #5 genome using BBMap (v38.79) (https://sourceforge.net/projects/bbmap/) with the flags minid=0.95 minaveragequality=30, and called variants with the BBTools scripts pileup.sh and callvariants.sh and the flags minreads=2 minquality=30 minscore=30 minavgmapq=20 minallelefraction=0.1.

##### Phylogenomics

A set of 56 universal single copy marker proteins (*56, 57*) was used to build a phylogenetic tree of the filament assemblies and related gammaproteobacterial genomes available in the IMG/M database (*58*). Marker proteins were identified with hmmsearch (version 3.1b2, hmmer.org) using a specific HMM for each of the markers. For every marker protein, alignments were built with MAFFT (v7.294b) (*59*) and subsequently trimmed with BMGE (*60*) using BLOSUM30. Single protein alignments were then concatenated and maximum likelihood phylogenies were inferred with iq-tree v2.0.3 (*61*) using the LG4X+F model. The tree was visualized in itol (*62*).

##### Secondary metabolism

Bacterial genomes were analyzed for secondary metabolite Biosynthetic Gene Clusters (BGCs) with antiSMASH v5.1.2 (*63*). Fungal BGC data for the fungus model system *Aspergillus nidulans* were retrieved from antismash-db. %BGC was calculated by summing the nucleotide length of each antiSMASH BGC region and dividing by total genome size.

##### Genome annotation and functional analysis

*Ca.* T. magnifica genomes were annotated using a JGI prokaryotic structural and functional genome annotation pipeline (https://img.jgi.doe.gov/docs/pipelineV5/) and loaded into IMG/MER database (*58*). Assignments of proteins to protein families, such as Pfam v.30 (*64*) and KEGG v.77.1 (*65*) in conjunction with the tools provided by IMG user interface were used to infer functional capabilities encoded by the genomes and to visualize chromosomal neighborhoods of genes of interest. Protein sequences of interest were exported from IMG/MER and alignments were built with MAFFT (v7.294b) (*59*).

### Supplementary Text

#### Evidence of thiotrophy in *Ca*. T. magnifica

Mangrove swamps accumulate fine sediment with high organic content. Under anoxic conditions sulfate reducing bacteria degrade organic matter producing large amounts of sulfide and sustaining sulfur oxidizing chemoautotrophic (thiotrophic) microbial communities. In Guadeloupean mangroves, sulfide produced by the reduced sediment ranges from 0.19 mM to 2.40 mM (*9*). It passively diffuses to the overlaying water where it gets rapidly oxidized by oxygen creating a steep redox gradient at the sediment/water interface. In order to characterize the microenvironment of *Ca.* T. magnifica, we brought to the laboratory filaments still attached to sunken leaves and immersed the leaves in a mesocosm-like setup on top of reduced mangrove sediment. Using microsensors we measured high and relatively stable concentrations of reduced sulfur species in the microenvironment of the filaments with concentrations fluctuating between 1.20 to 1.79 mM (Fig. S10) while no sulfide was detectable in the overlaying water.

Genomics and oxidoreductase activity experiments have shown that other *Thiomargarita* spp. can use reduced sulfur species as electron donors (*23, 24, 66*). Thiotrophic gammaproteobacteria from sulfidic environments are known to store elemental sulfur in membrane bound vesicles (*67, 68*). In order to determine if *Ca.* T. magnifica filaments display an active thiotrophic metabolism, we placed lightly fixed fully hydrated cells in an environmental scanning electron microscope and interrogated the presence of internal sulfur granules using Energy Dispersive X-ray Spectroscopy (EDXS). Images collected by secondary and backscattered electron detectors clearly showed the presence of bright round-shaped areas of on average 2.33 ± 0.46 µm in diameter (Fig. S10B and C). EDXS microanalysis (Fig. S10A) and sulfur mapping (Fig. S10F) clearly show that these areas are sulfur-rich and therefore correspond to sulfur globules (also visible on the TEM as numerous electron lucent vesicles 2.40 ± 1.03 µm in diameter).

#### Estimation of the genome copy number

We analyzed in 3D three datasets coming from three different cells after DNA was fluorescently labeled with DAPI. We visualized an entire 2.39 mm long cell (cell H, Table S2), as well as two cell portions of 195 µm (cell L, Table S2) and 660 µm (cell M, Table S2). After segmentation of the DNA clusters labeled with DAPI, we used the ORS Dragonfly© multi-ROIs analysis tool and counted 7647, 1518 and 3910 individual objects for the three cells respectively. The volume of individual DNA clusters ranged from 0.48 to 3511 um^3^. We assumed that the smallest DAPI stained DNA clusters, with a volume of 0.48 um^3^, represented a single genome copy and observed that larger clusters represented multiple chromosomes too close to one another to be segmented as individual objects. Therefore, we divided larger DNA clusters by the volume of the smallest ones (0.48 um^3^) and estimated the total number of genome copies for each data set. We then divided the total number of genome copies by the length of the cell analyzed allowing us to express the polyploidy level per millimeter of filament and extrapolate to a fully grown 20 mm long cell (see Table S2). We estimated that *Ca.* T. magnifica cells present on average 36,880 ± 7,956 genome copies per millimeter of filament and extrapolated that fully grown 20 mm cells would therefore present 737,598 ± 159,115 genome copies.

#### FISH investigations

The specificity of the FISH probe Thm482 was confirmed by the TestProbe tool from the SILVA database. In addition, we extracted all 16S rRNA gene sequences from the published *Thiomargarita* metagenome as well as from our five *Ca.* T. magnifica single-cell assemblies and further confirmed that the Thm482 probe only matched the *Thiomargarita* 16S rRNA sequence.

The FISH investigations were conducted on 4 different cells. From each cell, triplicate sections from 7 different locations consistently showed the same results, representative exemplars of which are shown in Figs. 2 and S6 to S8.

#### Extended description of *Ca.* T. magnifica metabolism based on the genome analysis

The large genomes of *Ca*. T. magnifica filaments encoded a wide range of metabolic capabilities, some unique and some shared with other large sulfur-oxidizing bacteria. A complete glycolysis pathway was present, which included a gene for glucose-6-phosphate isomerase; this gene was reported to be absent from previously sequenced single cells of *Ca*. T. nelsonii (Table S9) (*24*). The tricarboxylic acid cycle is also complete and can partially function in both directions due to the presence of 2-oxoglutarate dehydrogenase complex and 2-oxoglutarate-ferredoxin oxidoreductase, as well as succinate dehydrogenase and fumarate reductase. Multiple anaplerotic enzymes are also encoded (Table S9). *Ca*. T. magnifica genomes encoded RuBisCO form I (Fig.S12) and other enzymes for carbon fixation via the Calvin–Benson–Bassham (CBB) cycle. Similar to *Ca.* T. nelsonii, sedoheptulose-1,7-bisphophatase and the fructose-1,6-bisphosphatase were missing and may have been replaced by a polyphosphate-dependent 6-phosphofructokinase (*24*).

While *Ca.* T. nelsonii were found to perform denitrification or dissimilatory and assimilatory reduction of nitrate to ammonia, the only enzymes found in *Ca.* T. magnifica genomes were dissimilatory/respiratory nitrate reductases, Nar and Nap. Although there were genes encoding proteins similar to NorD and NorQ, accessory proteins of nitric oxide reductase, its catalytic subunits NorCB were missing, and so were genes encoding assimilatory nitrate reductase, both types of nitrite reductase (NirBD and NirS), and nitrous oxide reductase. Instead, a complete set of urease subunits was present (Table S9). This suggests that *Ca.* T. magnifica likely obtains ammonia for growth from organic sources, whereas nitrate is used exclusively as an electron acceptor.

Like other large sulfur-oxidizing bacteria, *Ca.* T. magnifica genomes encoded a set of genes for sulfur and hydrogen oxidation. Genes encoding two Ni-Fe hydrogenases with a complement of accessory proteins were found. Both sulfide-quinone-oxidoreductase Sqr and flavocytochrome c sulfide dehydrogenase FccAB were present, reverse dissimilatory sulfite reductase pathway, as well as thiosulfate-oxidizing sox genes. As in *Ca.* T. nelsonii, a group I catalytic intron was inserted into the *Ca.* T. magnifica *dsrA* gene encoding alpha subunit of dissimilatory sulfite reductase (*23*). *Ca.* T. magnifica genomes encoded a branched respiratory electron transport chain, which included NADH dehydrogenase, succinate dehydrogenase, ubiquinol-cytochrome c reductase, cbb3-type cytochrome c oxidase, and cytochrome d ubiquinol oxidase, as well as heterodisulfide reductase and sodium-transporting Rnf complex. The genomes also encoded both F-type and V-type ATPases.

In addition to the common bacterial secretion systems, a type VI secretion system (T6SS) was present in all *Ca.* T. magnifica genomes (Table S10), and some of its components were found in *Ca.* T. nelsonii. Often found in bacterial pathogens, this secretion system is analogous to contractile tails of bacteriophages and is known to mediate contact-dependent killing of neighboring cells via intracellular delivery of toxic effectors (*69*). T6SS consists of an inner tube composed of Hcp protein capped with VgrG and proline-alanine-alanine-arginine (PAAR) repeat-containing proteins surrounded by a sheath made of TssB/TssC heterodimers (also known as ImpB/ImpC or VipB/VipC). Other components of T6SS include a transmembrane complex composed of three subunits, TssJ/TssL/TssM (also known as ImpJ/ImpK/ImpL) and a baseplate, which is connected to the transmembrane complex and serves as a platform for inner tube and sheath polymerization. The injection process presumably starts with rearrangement of baseplate components leading to sheath contraction, opening of the baseplate and release of the inner tube and its payload into the target cell. Known T6SS effectors include peptidoglycan hydrolases, phospholipases, nucleases, ADP-ribosyltransferases, and pore-forming proteins (*70*); however, the vast majority of T6SS effectors remain unknown. Some of them can be identified based on the presence of VgrG or PAAR domain in their N-terminus, while others are characterized by the presence of other signature motifs, such as rearrangement hotspots (RHS), YD repeats, MIX motifs and FIX motifs (*71*). None of the known T6SS effectors were immediately recognizable in *Ca.* T. magnifica genomes; however, these genomes did encode between 8 and 22 RHS/YD repeat proteins (Table S11), as compared to 0 to 5 in closely related genomes. Three genes encoding RHS proteins were located near the T6SS component *vgrG*. The large variability of RHS protein count in *Ca.* T. magnifica may represent natural variation in the population or alternatively be an artifact of assembly due to the length and repetitive nature of these proteins. It is likely that T6SS found in *Ca.* Thiomargarita genomes confers a competitive advantage in the course of inter- and intraspecific conflict.

Among protein families, which have undergone significant expansion in *Ca.* T. magnifica, there were multiple families associated with genome rearrangements including mobile genetic elements, introns, and site-specific recombinases. *Ca.* T. nelsonii single cells possess the same genomic features (*23, 24*). In addition, *Ca.* T. magnifica genomes were riddled with a myriad of toxin/antitoxin systems (Table S11). While some of these systems may be involved in maintaining genetic material, such as mobile genetic elements, others may play regulatory roles in stress response, adaptation, as well as contribute to the complex lifestyle (*72*).

Expansion of other protein families have also been attributed to complex lifestyle and morphological changes: in addition to metacaspases (PF00656) described by Flood et al., 2016 (*23*), these include multiple copies of protein kinase domain (PF00069) and another family of peptidases and inactivated derivatives called CHAT (PF12770) (*73*). Curiously, nearly a third of caspase domains are associated with another highly overrepresented domain, called FGE-sulfatase (PF03781). Similarly, nearly a third of protein kinase domains are associated with FGE-sulfatase. This domain is known to generate formylglycine, which is the catalytic residue of sulfatases, and inactivation of a human formylglycine-generating enzyme SUMF1 leads to a multiple sulfatase deficiency (*74*). In bacteria characterized representatives of this family are iron(II)-dependent oxidoreductases EgtB and OvoA catalyzing C-S bond formation in the biosynthesis of ergothioneine and ovothiols (*75*). One of the FGE-sulfatase family proteins in *Ca.* T. magnifica is a likely bifunctional OvoA protein with 5-histidylcysteine sulfoxide synthase and mercaptohistidine methyltransferase activities. However, the function of more than a hundred copies of a gene encoding FGE-sulfatase in these genomes is unclear. A few copies of a gene encoding a predicted sulfatase (PF00884) were present in *Ca.* Thiomargarita genomes, but its abundance was not nearly enough to explain the expansion of the FGE-sulfatase family. Given its association with other domains implicated in complex life cycle and cell morphology, we hypothesize that the FGE-sulfatase family in *Ca.* Thiomargarita is also involved in these processes.

Another remarkable feature of *Ca.* T. magnifica was a significant rearrangement of the core genes involved in cell division and morphogenesis. Whereas all genes necessary for lipid-linked peptidoglycan monomer biosynthesis and export were present (Table S8), many core cell division proteins involved in Z ring assembly and regulation were missing. Of the main Z ring components, only a cytoskeletal protein FtsZ was found; protein FtsA, which tethers FtsZ to the membrane, and protein ZipA interacting with FtsZ and essential division peptidoglycan synthases were absent. FtsZ is part of well-conserved dcw (“division and cell wall”) operon, which in most other large sulfur-oxidizing bacteria also includes FtsA and a key late divisome protein FtsQ along with peptidoglycan monomer synthesis enzymes MurC, MurG, and Ddl. The overall structure of dcw operon was preserved in *Ca.* T. magnifica and *Ca.* T. nelsonii Bud S10, but FtsA and FtsQ were conspicuously missing (Fig. 3B). Consistent with the absence of FtsA, a widely conserved complex FtsE-FtsX, which plays many roles in promoting Z ring assembly including regulation of FtsA dynamics, was also missing from *Ca.* Thiomargarita genomes. In contrast, ZapA and ZapD proteins, which interact with FtsZ to promote Z ring assembly and stability, were present, and so was ZapA-interacting protein ZapB.

Even more remarkably, none of the late divisome proteins were found in *Ca.* Thiomargarita genomes. These include peptidoglycan polymerizing glycosyltransferase FtsW, peptidoglycan transpeptidase FtsI (peptidoglycan-binding protein 3), a key divisome complex FtsQLB, which recruits and regulates peptidoglycan synthesis activities, and FtsK protein, which bridges Z ring with late divisome components (*26*). We cannot exclude the possibility that the lack of these proteins was due to the assembly gaps, but the fact that not a single one of them was found in any of *Ca.* T. magnifica or *Ca.* T. nelsonii genomes suggests that they are truly missing and not merely an artifact of incomplete sequences.

While the complement of divisome proteins encoded by *Ca.* T. magnifica genomes was greatly reduced, components of cell elongation complex have been duplicated, which likely happened in the ancestor of *Thiomargarita* spp (Fig. S13-15). Elongasome components, which include cytoskeletal protein MreB required for maintenance of cylindrical cell shape, its interacting partners MreC, MreD and regulator of polymerization RodZ (Fig. S13), as well as peptidoglycan polymerizing glycosyltransferase RodA and peptidoglycan transpeptidase MrdA (peptidoglycan-binding protein 2), were organized in *Ca.* Thiomargarita into 2 chromosomal clusters. The first included MreBCD proteins, RodA and MrdA, as well as membrane-bound lytic murein transglycosylase MltB and rare lipoprotein A (Fig. 3B). The second included RodZ and genes unrelated to elongasome, such as 4-hydroxy-3-methylbut-2-enyl-diphosphate synthase ispG (Fig. 3B). Comparison of *mre* and *rodZ* regions of *Ca.* Thiomargarita to those of other large sulfur-oxidizing bacteria revealed duplications of *rodZ*, *mreD* and *mrdA*, with both copies adjacent on the chromosome. While the physiological effect of this increase of gene dosage is unclear, it is likely related to the complex morphology and life cycle of *Ca.* T. magnifica. Alternatively, they may compensate for the lack of peptidoglycan synthesis components of the divisome.

#### Developmental cycle of *Ca.* T. magnifica

Because we observed fully grown filaments releasing their most apical segment live in the lab, we hypothesized that the released terminal segments represent a dispersive stage of the developmental cycle (Fig. 1C, S1). To test this hypothesis, we quantified the biovolume of the apical buds and compared it to the biovolume of a small filament (since the bacterium is uncultured). The smallest filament analyzed was observed with hard x-ray tomography and had a volume of 2.37×10^-13^ m^3^ which corresponds almost exactly to the volume of terminal buds from the apical pole of the fully-grown filament (2.1 and 2.4×10^-13^ m^3^). It is likely that the rod-shaped terminal segment of filaments represents dispersive daughter cells which get released to the environment and eventually attach and grow on a new substrate. The newly attached daughter cell apparently first reshapes itself into a thin filamentous cell while conserving the same volume. Young filaments then grow in the vertical direction with the apical pole starting to constrict to form new buds (Fig. 1C, S1).

Based on field observations (by O. Gros, co-author), the *Ca.* T. magnifica developmental cycle may be somewhat analogous to *Zoothamnium niveum,* a giant colonial ciliate that sometimes co-occurs on the same substrate. Like *Ca*. T. magnifica, elongated colonies of *Z. niveum* grow up to 1.5 cm high, emerging above the competitive biofilm to provide an optimal access to hydrogen sulfide and oxygen to its sulfur-oxidizing gammaproteobacterial symbiont (*30*). Like *Ca.* T. magnifica, specialized cells eventually detach from the colony, disperse, settle in favorable environments, and grow into colonies. We observed co-occurring *Ca.* T. magnifica cells and *Z. niveum*, sometimes on the same decaying leaf. The gigantism and life cycle of *Ca.* T. magnifica may therefore allow them to exploit a niche so far known to be occupied only by Eukaryotes such as *Z. niveum.* Just like the sulfur-oxidizing ectosymbionts escape the biofilm competition by growing vertically through their association with the ciliate, *Ca.* T. magnifica grows above the biofilm where it is still exposed to high sulfide concentrations (Fig. S11).

**Fig. S1.**
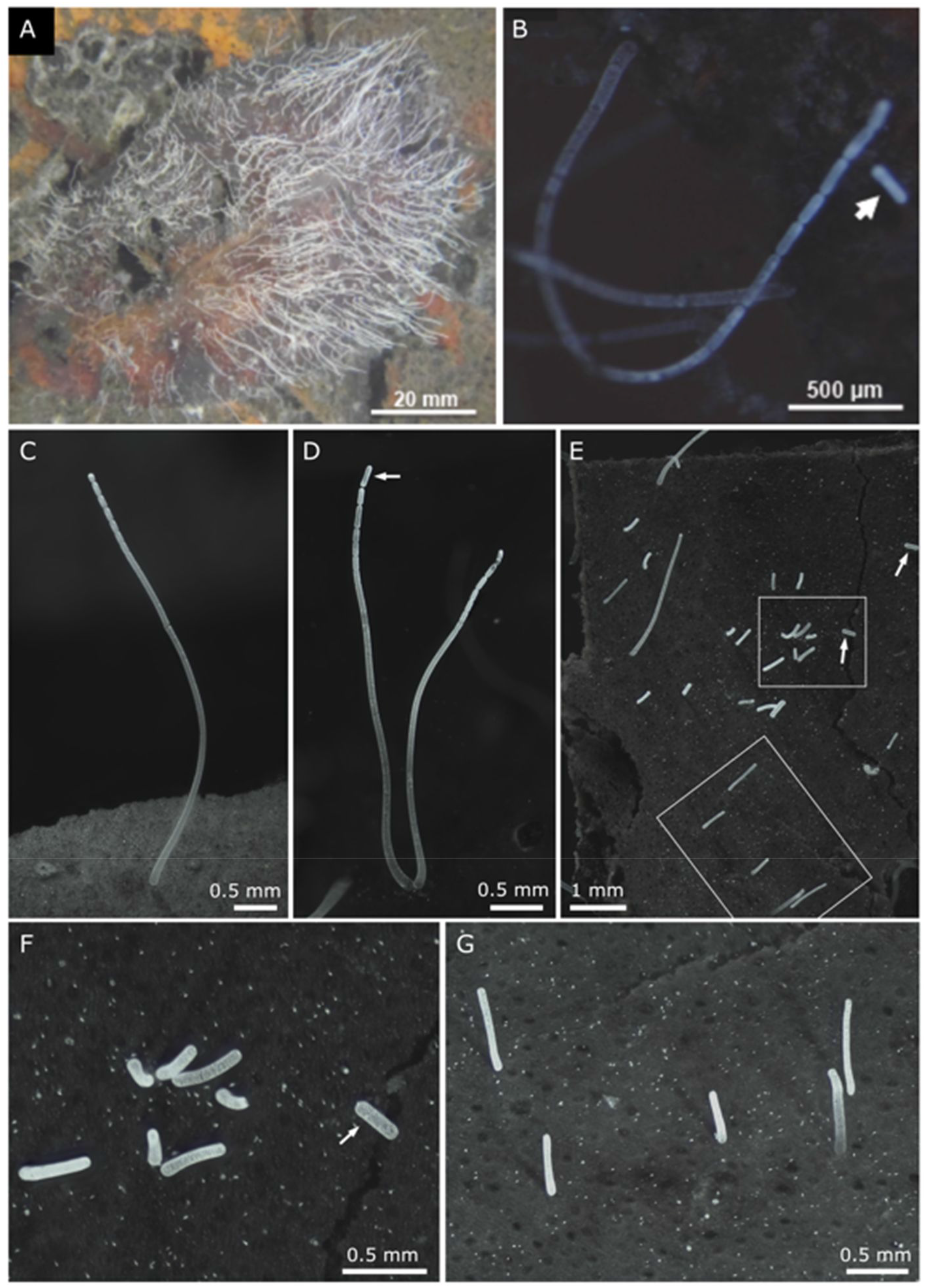
Light micrographs of *Ca*. Thiomargarita magnifica attached to sunken leaves of *Rhizophora mangle*. A. Hundreds of filaments develop on sunken dead leaves partially buried in the sulfidic sediment of the mangrove. Similar patches were occasionally observed growing on partially buried plastic and oyster shell debris. The cells appeared bright white due to their intracellular elemental sulfur stores. B-D. Detail of budding cells showing a gradual constriction at the filament apex. The terminal segment (arrow) is still attached to the mother cell on D and has just been released into the water column for dispersion on B. E. Area of a sunken leaf with recently released terminal segments (arrows) and newly settled filaments of various sizes. Note that these young filaments (< 3 mm) are not displaying apical constrictions yet. The two rectangles’ areas are shown at higher magnification on F and G.

**Fig. S2.**
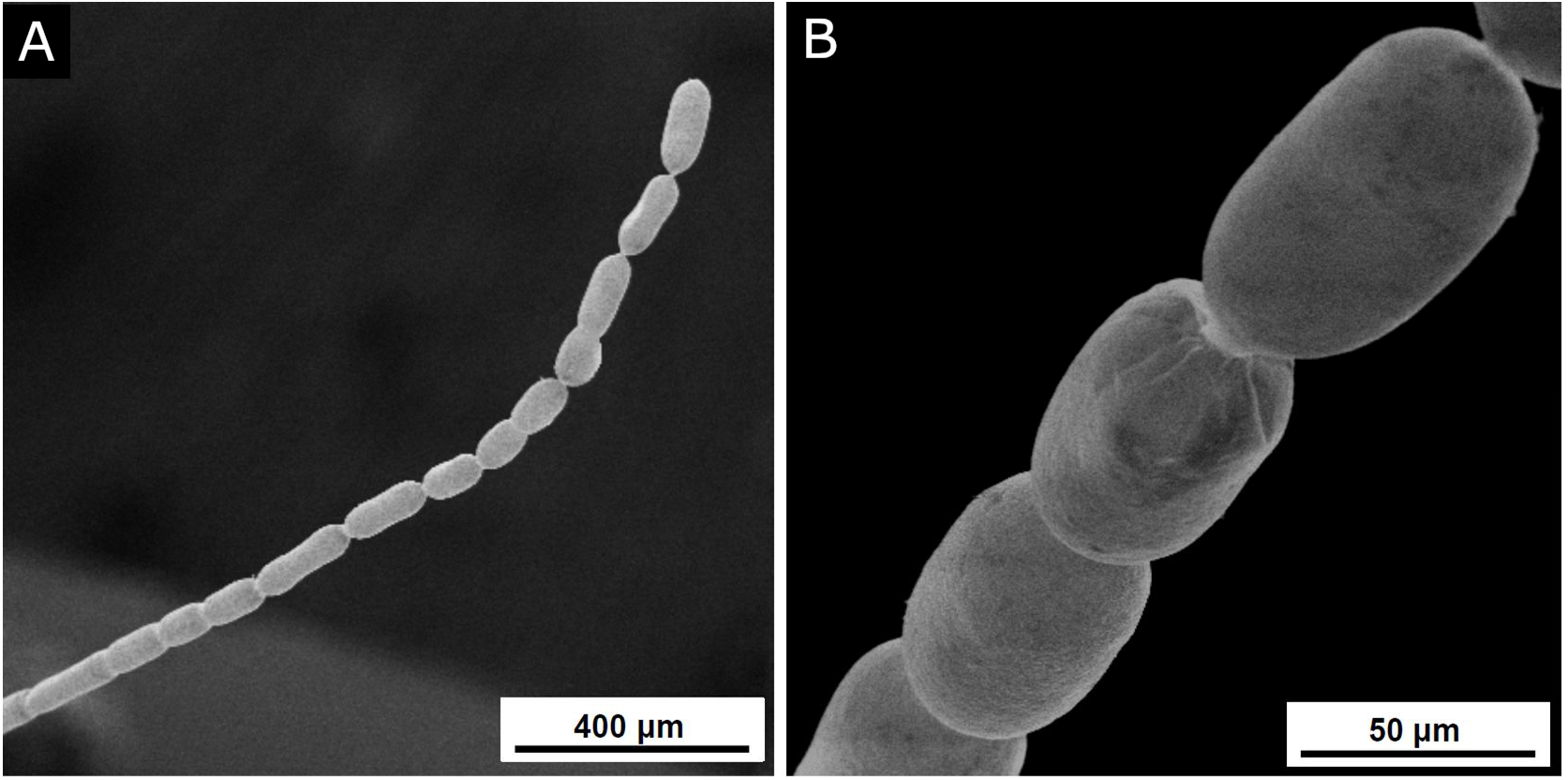
Scanning electron microscopy observation of an individual *Ca*. Thiomargarita magnifica. A. Detail of the last 2 mm of the cell apex characterized by multiple budding daughter cells. B. Higher magnification showing the smooth surface of the cell wall and the absence of epibiotic bacteria or biofilm.

**Fig. S3.**
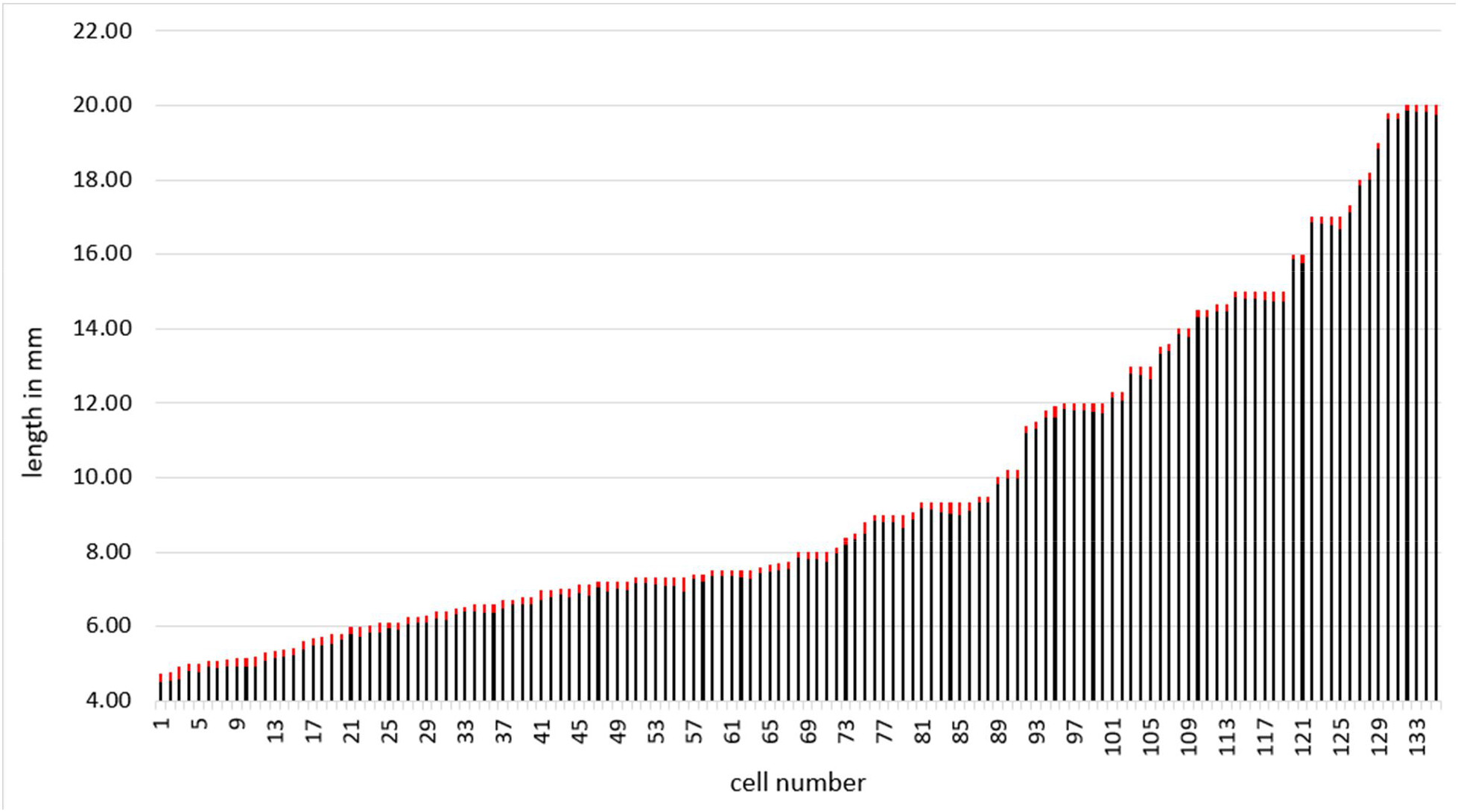
Lengths of 135 *Ca.* Thiomargarita magnifica budding cells measured under a stereomicroscope. The smallest filament observed with a terminal segment here was 4.73 mm and the largest filaments were 20 mm. The length of the most apical segment is displayed in red.

**Fig. S4.**
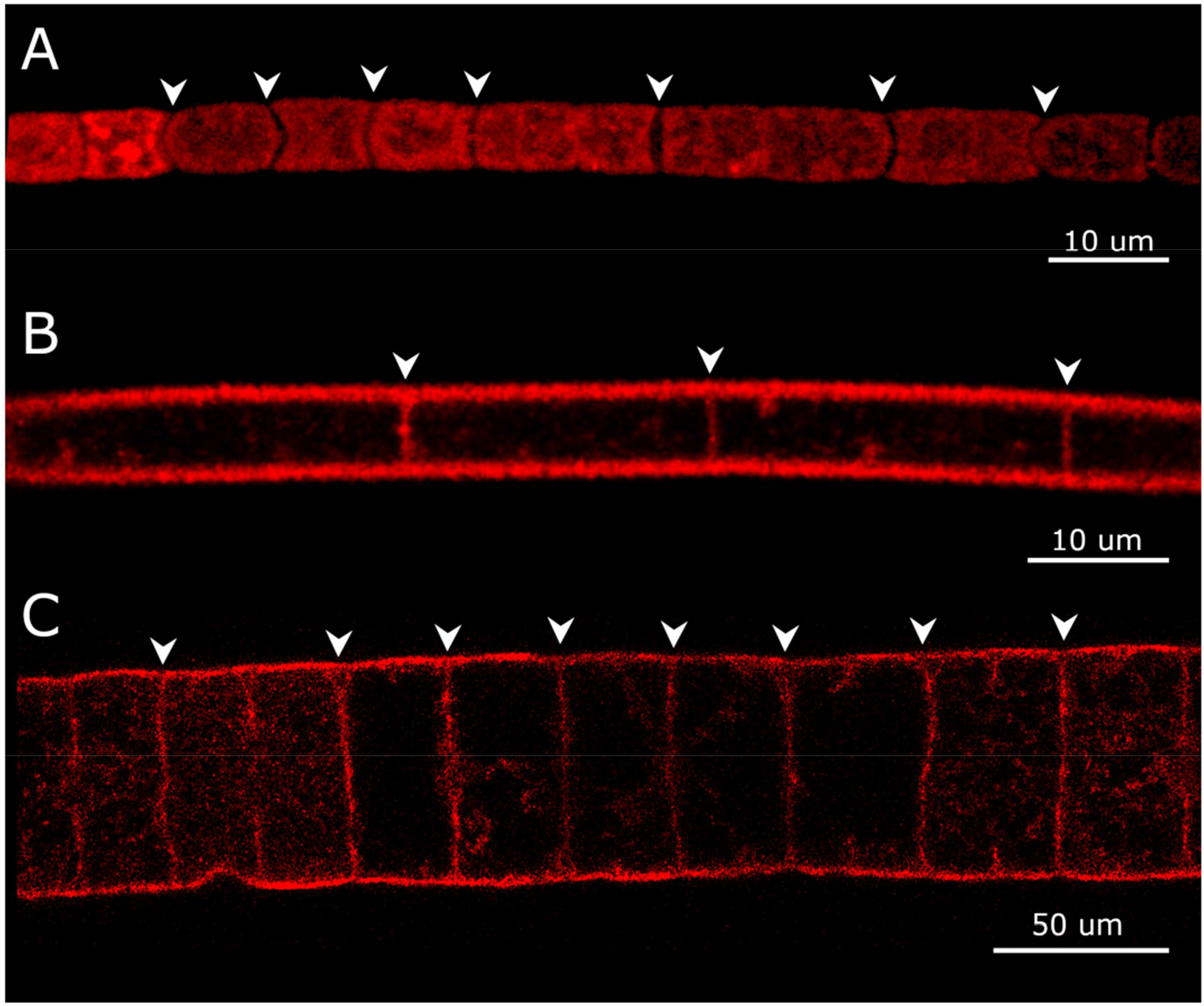
Confocal laser scanning microscopy observations of three multicellular large filamentous bacteria after fluorescent labeling of membranes using FM 1-43x dye. A. The separations between the cells (white arrowheads) of the Cyanobacterium *Microcoleus vaginatus* are clearly visible after FM 1-43x staining. B. The membrane septum separating the vacuolated cells of a *Beggiatoa*-like filament are clearly stained by the membrane dye. C. The membrane septum separating the large vacuolated cells of the *Marithrix*-like bacterial filaments are clearly stained by the membrane dye.

**Fig. S5.**
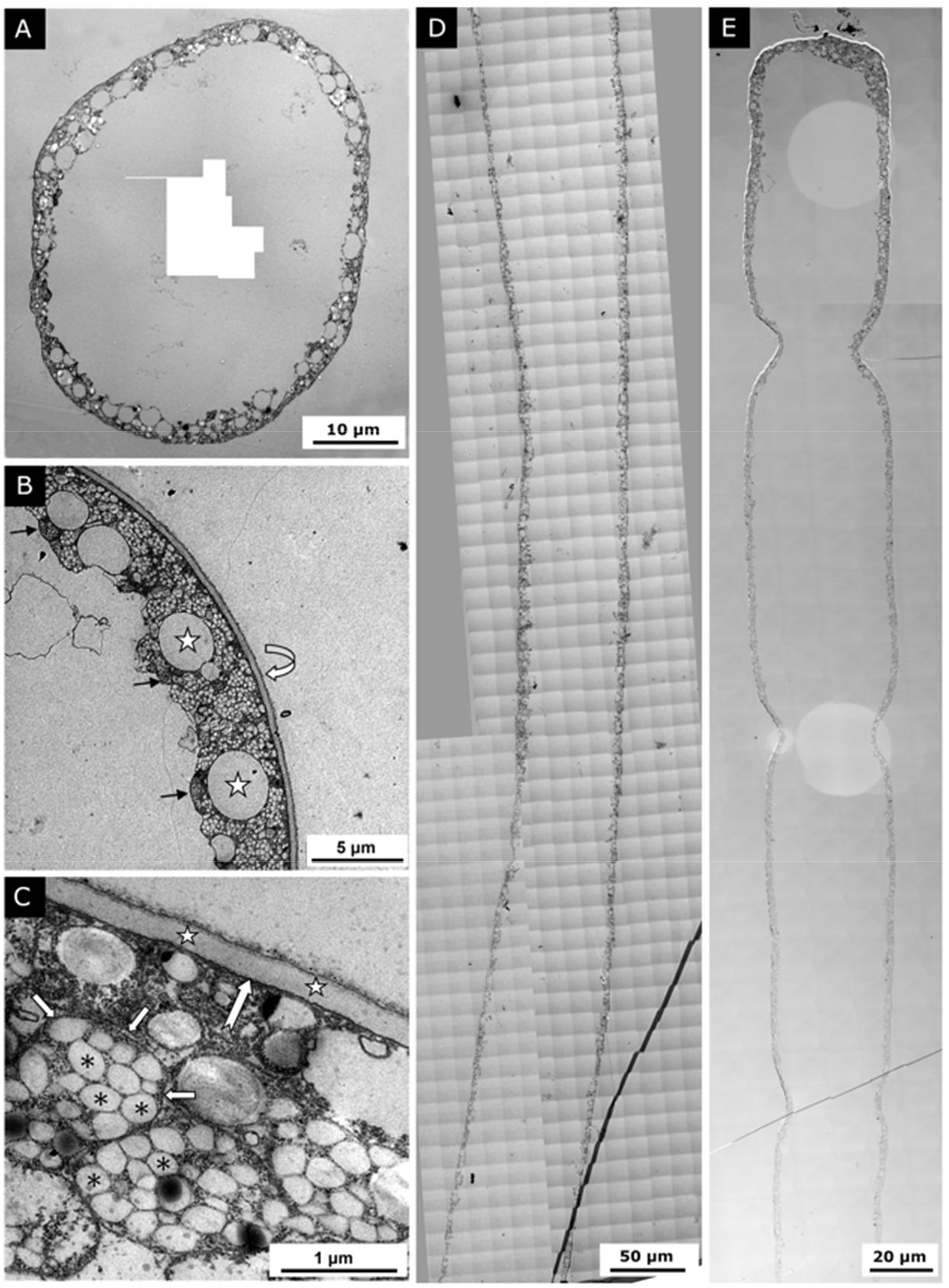
Transmission electron microscopy observation of *Ca.* T. magnifica. A. Photo montage of a cell cross section. The cytoplasm is organized at the periphery of the cell around a large central vacuole. B. The cell envelope is represented by a thick outer layer covering the cytoplasmic membrane (curved white arrow). Electron lucent vesicles correspond to elemental sulfur granules (stars) dissolved by ethanol during sample dehydration. Pepins are located adjacent to the central vacuole (black arrows). C. Detailed view of the cytoplasmic membrane (big arrow) covered by a cell wall (stars). The cytoplasm is filled with organelles, each delimited by a membrane (small arrows), that sometimes contain multiple electron lucent vesicles (asterisks). D. Photomontage (764 images taken at magnification 3000x) of a cell longitudinal section 850.6 µm long. The section corresponds to the middle part of the filament. No septum was observed and the cytoplasm, cell membrane and central vacuole appear continuous throughout the whole section. E. Photomontage (349 images taken at magnification 1400x) of a cell longitudinal section 374 µm long. The montage shows the continuity of the cell and the absence of any membrane septum, even at the constriction sites.

**Fig. S6.**
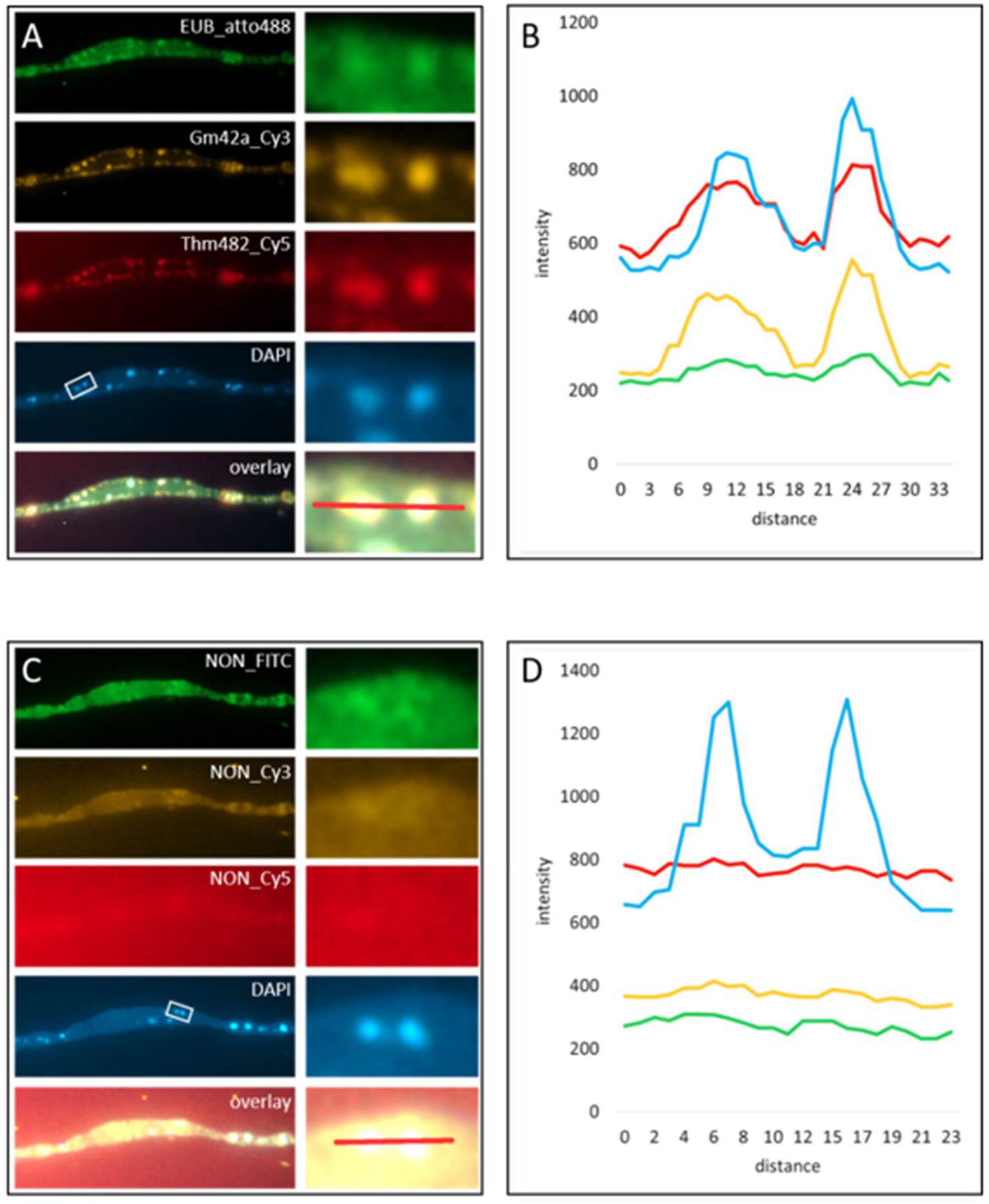
Fluorescence *In Situ* Hybridization (FISH) with general, specific and nonsense probes and DNA staining with DAPI on sections of resin-embedded *Ca.* Thiomargarita magnifica cells. A. Pepins are labeled with the general bacterial probe (green), a gammaproteobacteria-specific probe (yellow), the *Thiomargarita* specific probe (red), or with DAPI (blue). The detail of the two pepins in the white rectangle on the DAPI image is shown at higher magnification on the right. B. The intensity profiles of a line crossing the two pepins from the insert is showing peaks in all four channels. C. FISH performed on sections of the same cell placed on the same slide show no hybridization of the nonsense probe. D. The intensity profiles of the line crossing the two pepins from the insert on B show a peak for the DAPI signal but no signal in any of the probe colors.

**Fig. S7.**
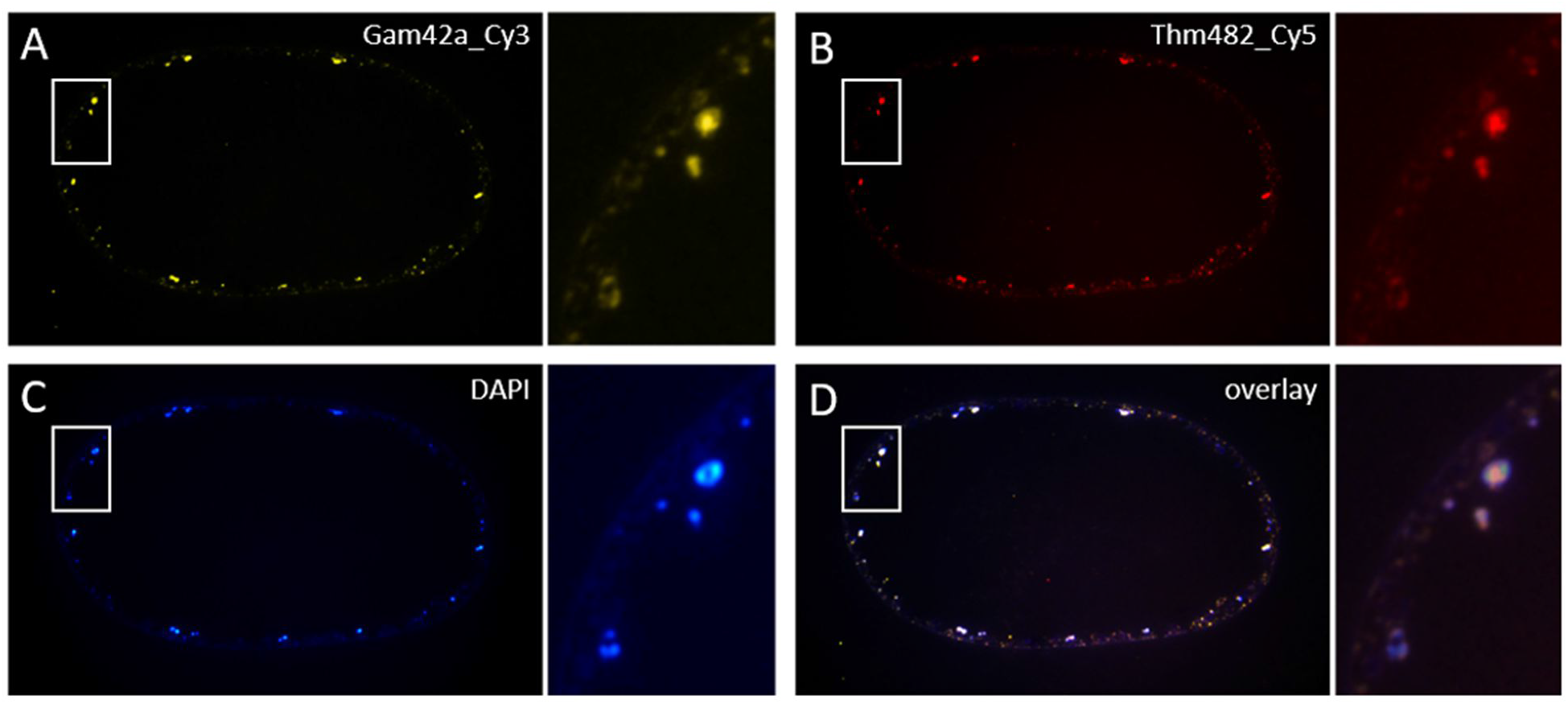
Fluorescence *In Situ* Hybridization (FISH) with gammaproteobacteria (A) and *Thiomargarita* (B) probes and DNA staining with DAPI (C) on a cross section of the stalk area of a *Ca.* Thiomargarita magnifica cell. The pepins are labeled with both gammaproteobacteria and *Thiomargarita* probes - *Thiomargarita* is a genus within the gammaproteobacteria - as well as by DAPI as shown by the overlay of the three images (D). A higher magnification of the area in the white rectangle is provided on the right of each image. It shows that while the FISH and DAPI signals colocalize in the pepins, they do not display the same pattern. On the larger pepin for instance, the ribosomes’ RNAs labeled in yellow and red are concentrated in the center of the organelle while the DNA labeled in blue shows higher signal on the sides.

**Fig. S8.**
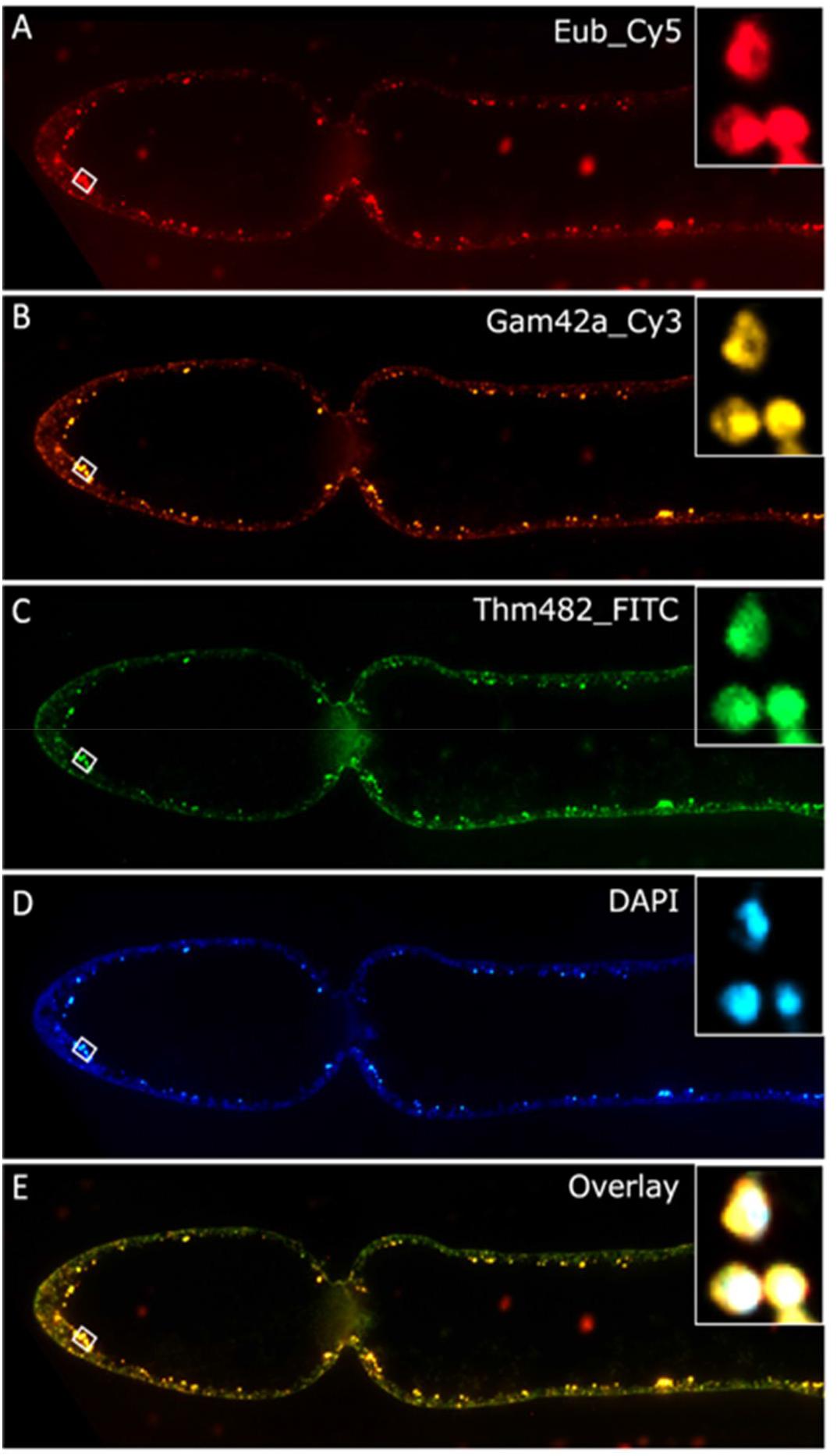
Fluorescence *In Situ* Hybridization (FISH) with probes targeting general bacteria (A, probe EUB labeled with Cy5); gammaproteobacteria (B, Gam42a probe labeled with Cy3); and *Thiomargarita,* a genus within gammaproteobacteria (C, probe Thm482 labelled with FITC); and DNA staining with DAPI (D); on a longitudinal section of the apical end of a *Ca.* Thiomargarita magnifica cell. On this longitudinal section pepins are present in the cytoplasm of the terminal segment as well as in the stalk before the constriction. Note that the terminal segment, like the rest of cell, presents a large central vacuole which occupies most of the cell volume. The pepins labeled with all three probes and DAPI are shown by the overlay (E). The detail of the three pepins in the white rectangle is given in the top right insert. Note that no epibiotic bacteria are detected with FISH.

**Fig. S9.**
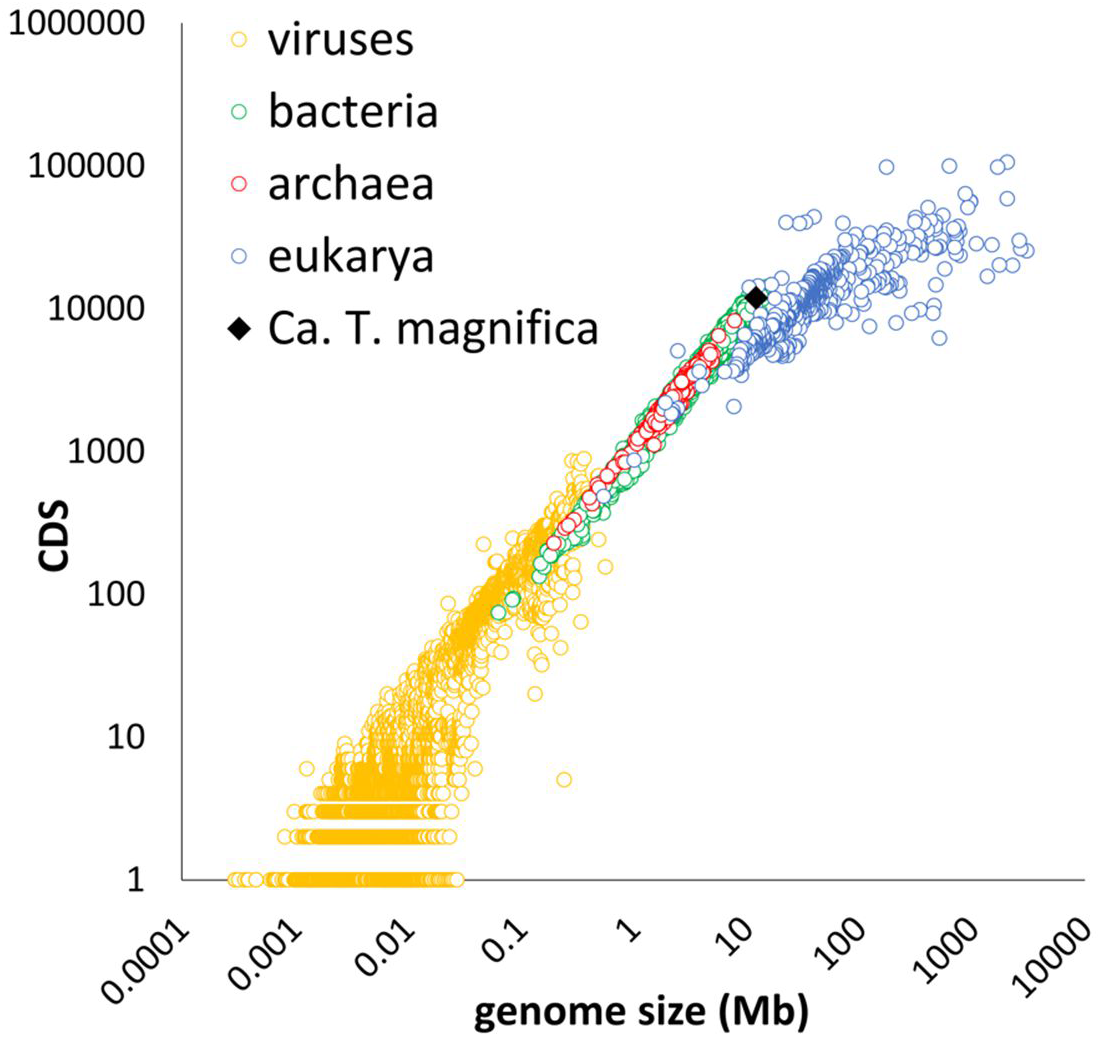
Total number of coding sequences (CDS) compared to the total genome size for all genomes retrieved from IMG/M. *Ca.* Thiomargarita magnifica’s genome is among the largest bacterial genomes with one of the highest number of coding sequences.

**Fig. S10.**
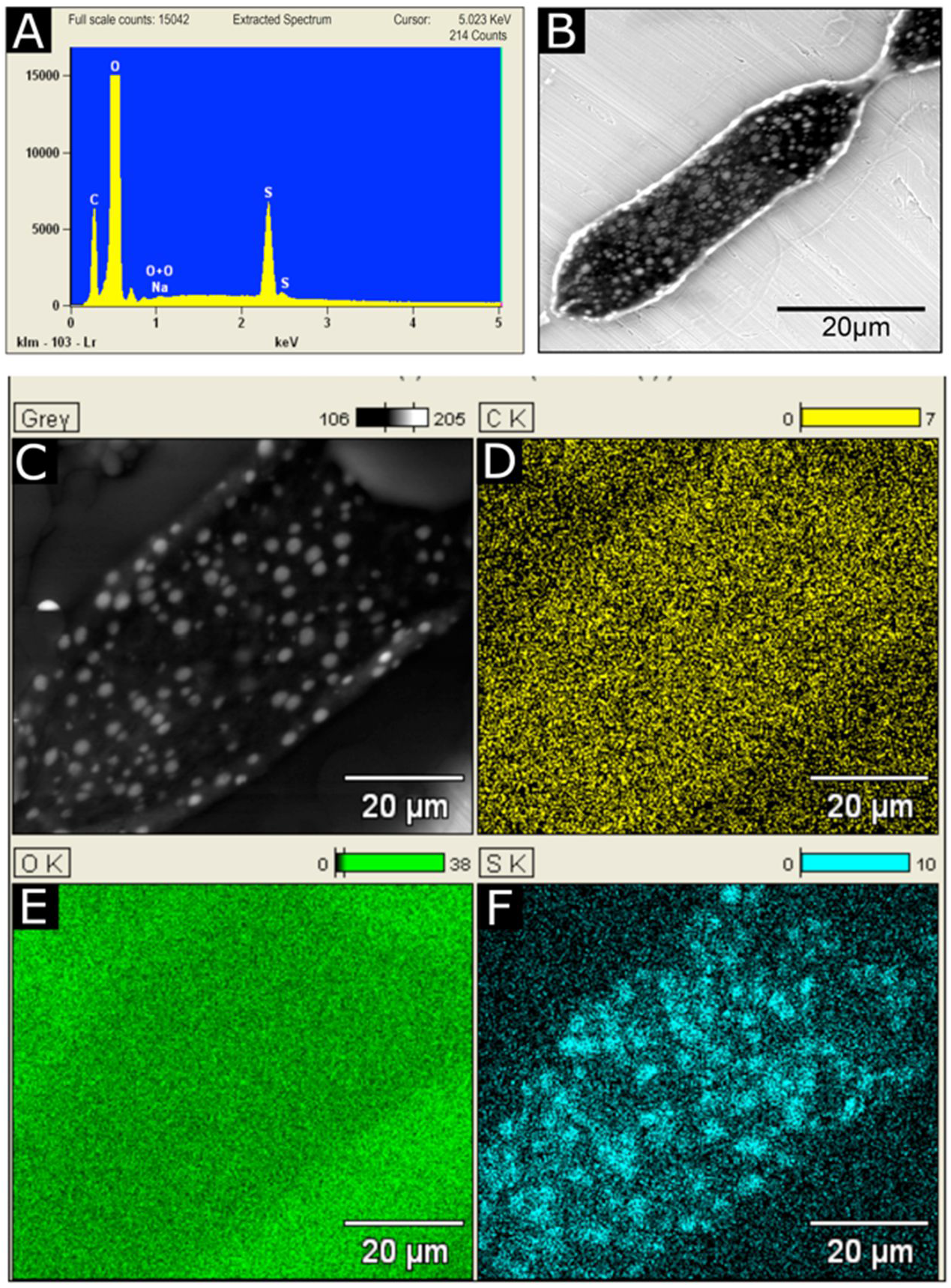
Environmental Scanning Electron Microscopy images and analyses recorded on slightly fixed *Ca.* Thiomargarita magnifica under 650 mPa water vapor atmosphere. A: EDXS spectrum acquired on the area corresponding to Figure S10B. Sulfur is clearly identified. B: Image collected with secondary electron detector with 15 kV accelerating voltage. The image is dominated by backscattered electrons due to the high penetration power of the incident electrons in the specimen. The resulting atomic number (Z) contrast reveals high Z number granules (sulfur) as bright areas embedded in the light (C, N, O, H) matrix in black. Secondary contrast is present but only visible at the periphery of the cell. C: Electron back scattered images obtained at 15 kV highlighting the sulfur granules as in B. Each empty granule appears white in a back-scattered electron image due to the intensity of the BSE signal which is strongly related to the atomic number of the chemical element. D-F: X-ray maps characterizing distributions of carbon (D), oxygen (E) and sulfur (F) in the sample. The sulfur map (F) clearly confirms that the bright areas appearing in the BSE images correspond to sulfur locations.

**Fig. S11.**
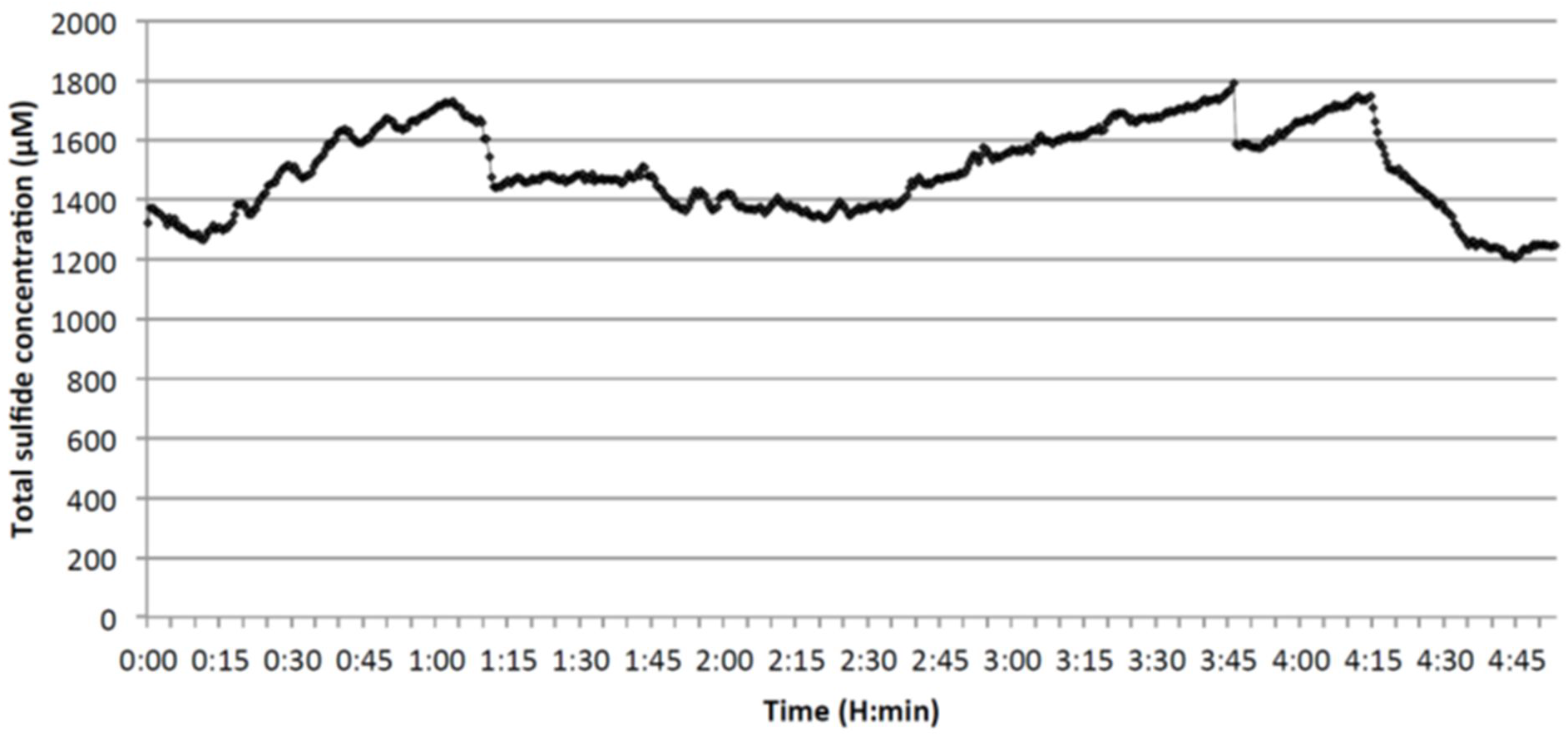
Variation of total sulfide concentration measured in the middle of a “bouquet” of *Ca*. Thiomargarita magnifica attached to sunken leaves of *Rhizophora mangle* over 5 hours.

**Fig. S12.**
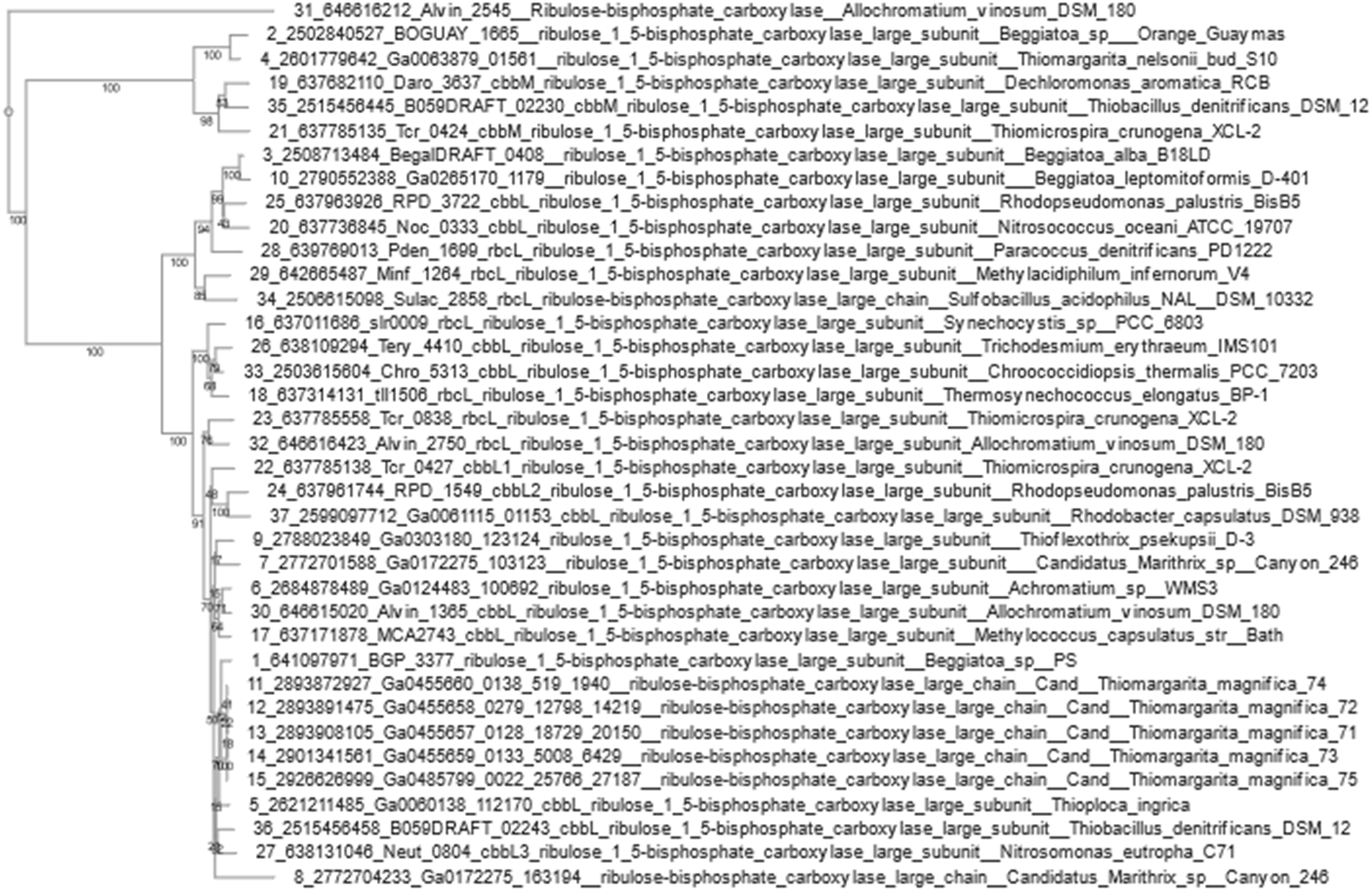
Phylogenetic tree of RuBisCo large subunit proteins encoded by *Ca*. T. magnifica and representative sequences from other genomes. *Ca*. T. magnifica rbcL sequences cluster within the type I clade, separately from type II RuBisCO encoded by *Ca*. T. nelsonii bud S10 genome.

**Fig. S13.**
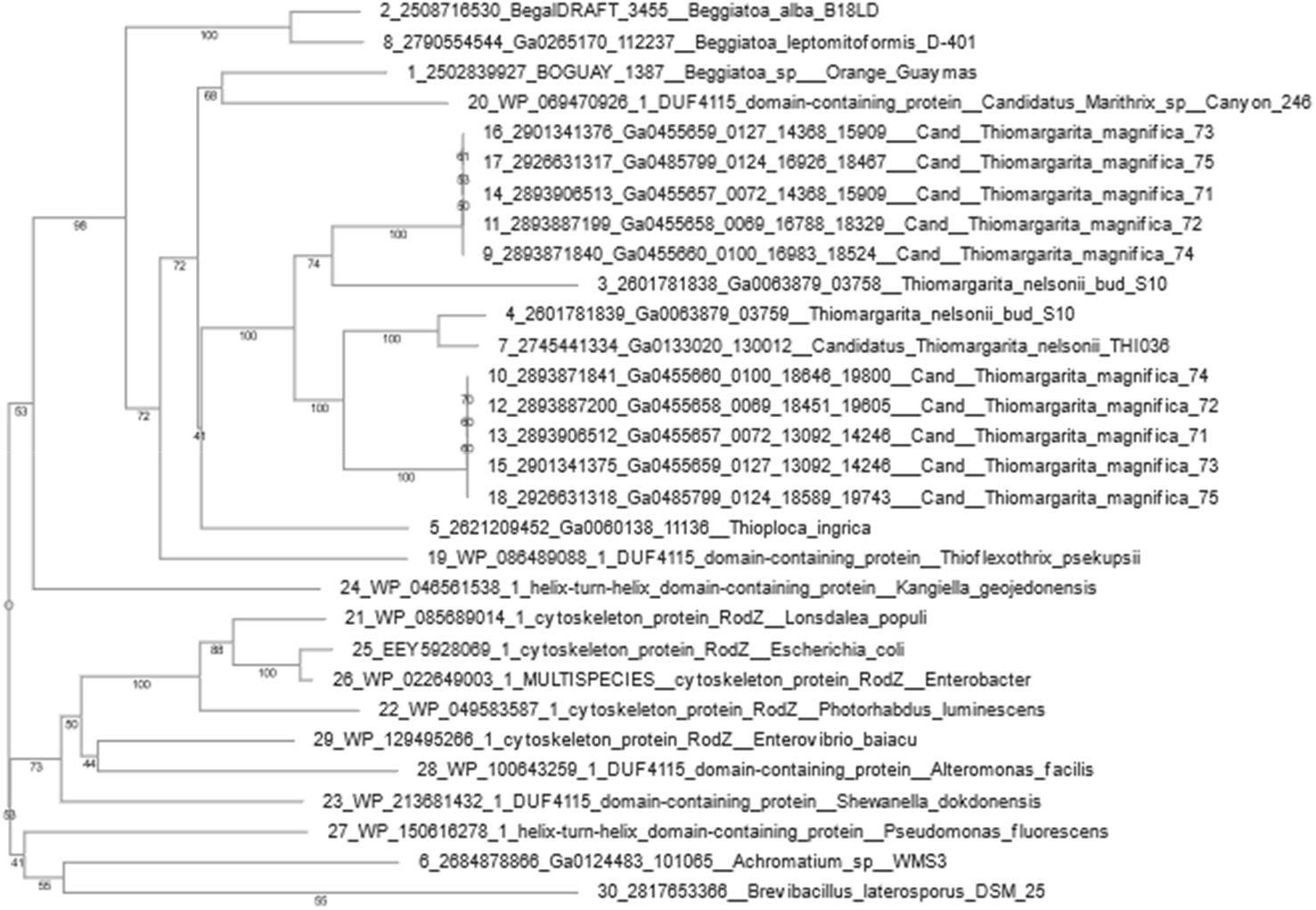
Phylogenetic tree of RodZ proteins encoded by *Ca*. T. magnifica draft genomes and selected reference sequences. Two copies of *rodZ* found next to each other in *Ca*. T. magnifica and *Ca*. T. nelsonii bud S10 are more similar to each other than to any other sequence, suggesting a duplication in a common ancestor of *Ca*. Thiomargarita spp.

**Fig. S14.**
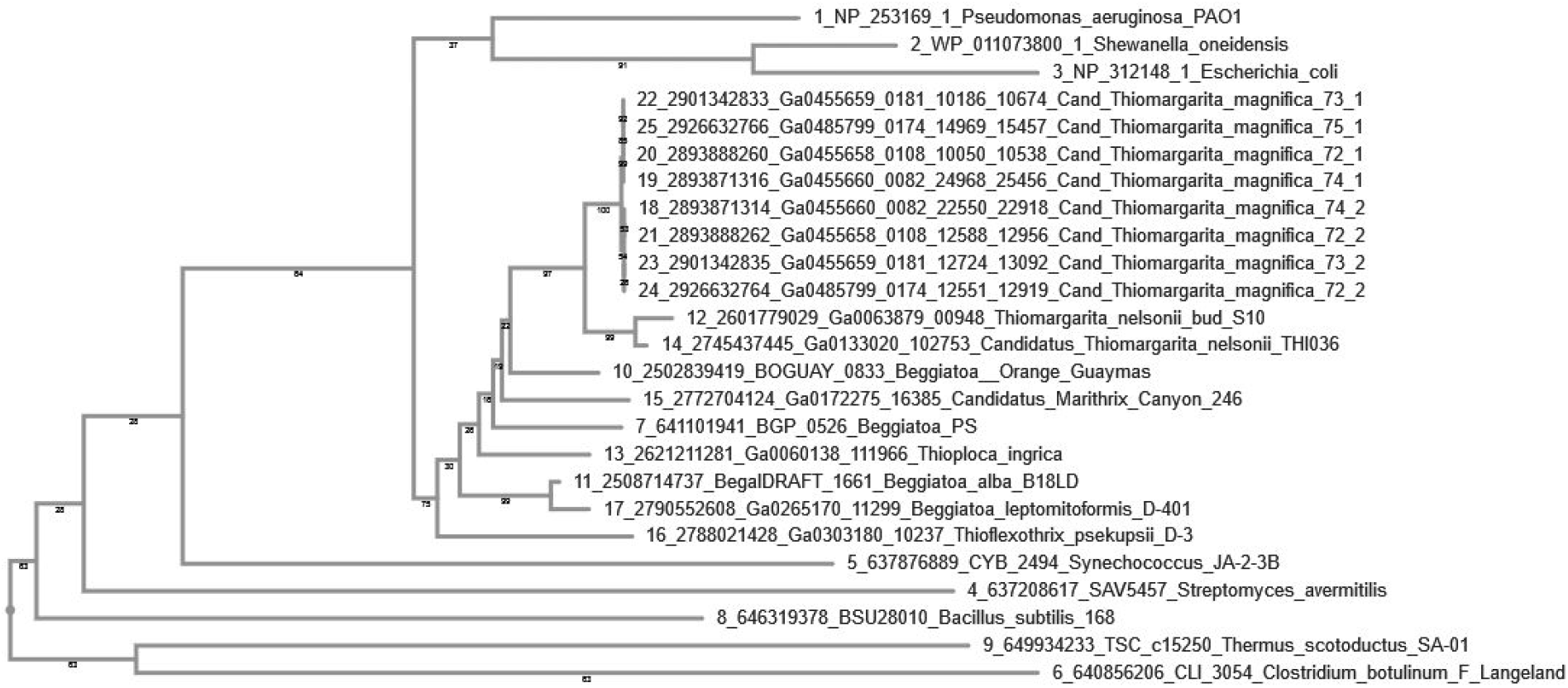
Phylogenetic tree of MreD proteins encoded by *Ca*. T. magnifica draft genomes and selected reference sequences. Two copies of the *mreD* gene found close to each other in *Ca*. T. magnifica are more similar to each other than to any other sequence, suggesting a duplication in *Ca*. T. magnifica.

**Fig. S15.**
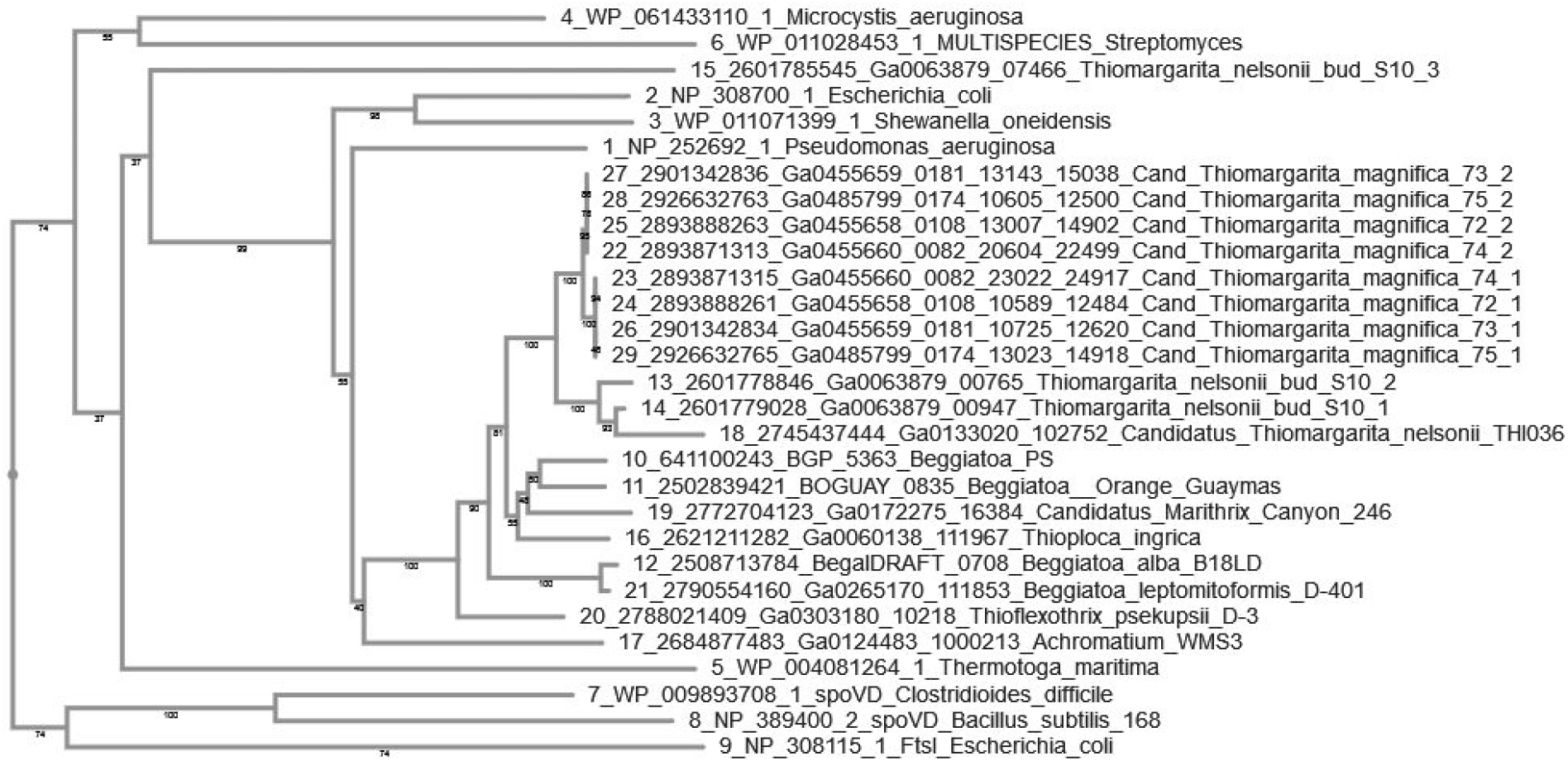
Phylogenetic tree of MrdA proteins encoded by *Ca*. T. magnifica draft genomes and selected reference sequences. Two copies of the *mrdA* gene found close to each other in *Ca*. T. magnifica are more similar to each other than to any other sequence suggesting a duplication in *Ca*. T. magnifica.

**Fig. S16.**
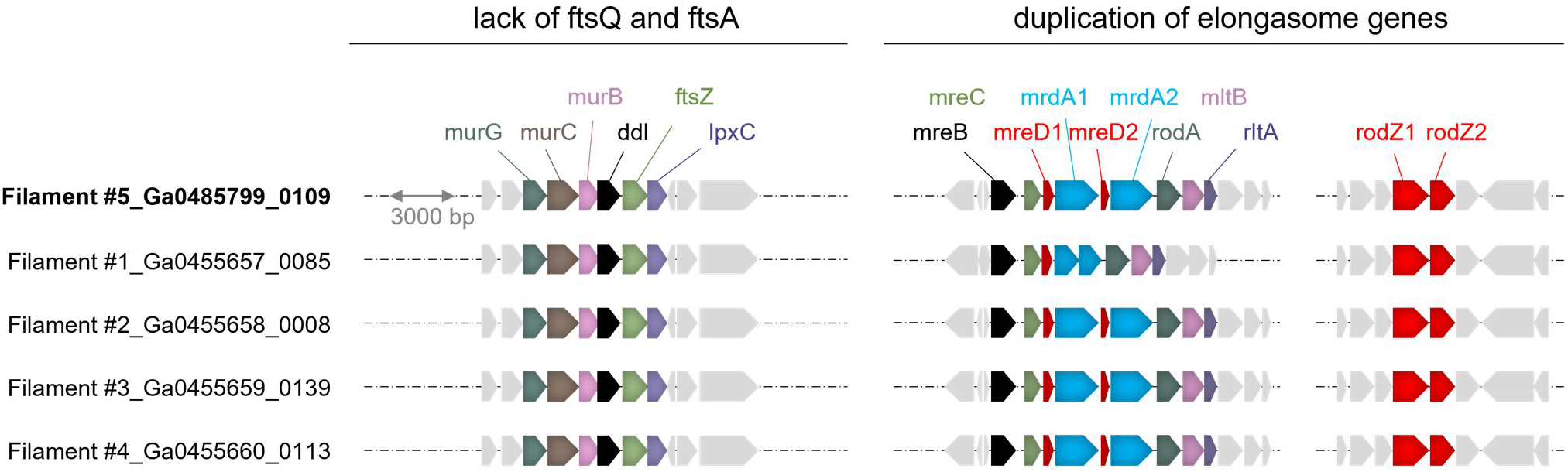
Detail of the *ddl*, *mreB* and *rodZ* gene neighborhood for the five *Ca*. T. magnifica draft genomes (see Fig. 3B).

**Table S1.** Summary of hard x-ray computed tomography scans parameters.

|  | cells A, B,<br>C, E & N | cell D |
| --- | --- | --- |
| scan date | 8/30/2019 | 8/27/2019 |
| total number of cells in sample | 5 | 1 |
| number of entire cells in the scan | 3 | 1 |
| dataset width (um) | 1755 | 984 |
| dataset height (um) | 1843 | 1008 |
| dataset depth (um) | 4668 | 6270 |
| number of tiles | 3 | 8 |
| pixel size (um) | 1.00 | 1.00 |
| number of projections per tile | 1601 | 801 |
| camera binning | 1 | 2 |
| source filter | air | air |
| source setting (kV) | 40 | 40 |
| source setting (uA) | 74 | 74 |
| source-RA distance (mm) | -9.2 | 0.0 |
| optical magnification | 4.0 | 4.0 |
| exposure time (s) | 15 | 10 |

**Table S2.** Summary of the *Ca.* T. magnifica observations with Hard X-ray Tomography (HXT) and Confocal Laser Scanning Microscopy (CLSM). A total of 14 cells were analyzed in 3D, ten of which were analyzed in their entire length. The table provides the detail of the length of the observed cell. When applicable the table also shows the cell minimum and maximum diameters, its total volume, cytoplasm volume and central vacuole volume as well as the percentage of vacuole volume relative to the whole cell. For three cells, the numbers of genome copies were estimated. This estimation is also provided as a number of genome copies per millimeter of cell and as an extrapolation for a fully grown 2 cm cell. Finally, the lateral resolution of the image as well as the z resolution and the number of tiles assembled in the dataset is provided.

| cell ID | analyzed with | entire cell | cell length (mm) | min diam (μm) | max diam (μm) | total cell volume (m<sup>3</sup>) | cytoplasm volume (m<sup>3</sup>) | vacuole volume (m<sup>3</sup>) | % of vacuole | est. genome copies | est. genome copies/mm | est. genome copies in 2cm cell | resolutio X - Y (μm) | resolution Z (μm) | tiles no. |
| --- | --- | --- | --- | --- | --- | --- | --- | --- | --- | --- | --- | --- | --- | --- | --- |
| A | HXT | Y | 1.19 | 17 | 33 | 2.37E-13 | - | - | - | - | - | - | 1 | 1 | 1 |
| B | HXT | Y | 1.54 | 17 | 29 | 3.50E-13 | - | - | - | - | - | - | 1 | 1 | 1 |
| C | HXT | N | 1.66 | 70 | 119 | - | - | - | - | - | - | - | 1 | 1 | 1 |
| D | HXT | Y | 6.37 | 43 | 83 | 1.42E-11 | 5.30E-12 | 8.89E-12 | 63 | - | - | - | 1 | 1 | 8 |
| E | HXT | Y | 9.66 | 47 | 96 | - | - | - | - | - | - | - | 1 | 1 | 3 |
| F | CLSM | Y | 1.92 | 54 | 96 | 6.59E-12 | 1.40E-12 | 5.19E-12 | 79 | - | - | - | 0.42 | 2 | 12 |
| G | CLSM | Y | 4.27 | 61 | 147 | 2.20E-11 | 5.91E-12 | 1.61E-11 | 73 | - | - | - | 0.63 | 2.73 | 5 |
| H | CLSM | Y | 2.39 | 39 | 62 | 4.29E-12 | 9.26E-13 | 3.36E-12 | 78 | 72419 | 30263 | 605256 | 0.23 | 1 | 14 |
| I | CLSM | Y | 3.35 | 66 | 112 | - | - | - | - | - | - | - | 1.27 | 2.73 | 6 |
| J | CLSM | Y | 2.32 | 52 | 87 | - | - | - | - | - | - | - | 1.24 | 2.73 | 3 |
| K | CLSM | Y | 2.83 | 67 | 71 | - | - | - | - | - | - | - | 0.92 | 2.95 | 8 |
| L | CLSM | N | 0.20 | - | - | - | - | - | - | 8931 | 45707 | 914140 | 0.1 | 1 | 1 |
| M | CLSM | N | 0.66 | - | - | - | - | - | - | 22882 | 34670 | 693397 | 0.23 | 1 | 1 |
| N | HXT | N | 1.97 | 58 | 101 | - | - | - | - | - | - | - | 1 | 1 | 2 |

**Table S3.** Reads, assembly and binning statistics for the five single amplified genomes of *Ca*. Thiomargarita magnifica.

|  | Filament 1 | Filament 2 | Filament 3 | Filament 4 | Filament 5 |
| --- | --- | --- | --- | --- | --- |
| Library total bases (Mb) | 816.5 | 1007.5 | 1659.2 | 1644.6 | 3057.3 |
| Read counts (x10<sup>6</sup>) | 5.5 | 6.7 | 11.1 | 11.0 | 20.4 |
| Assembly size (Mb) | 18.87 | 16.88 | 42.45 | 16.26 | 16.48 |
| Number of contigs | 6613 | 4720 | 27356 | 2383 | 2196 |
| Number of bins | 3 | 2 | 6 | 2 | 2 |

**Table S4.**
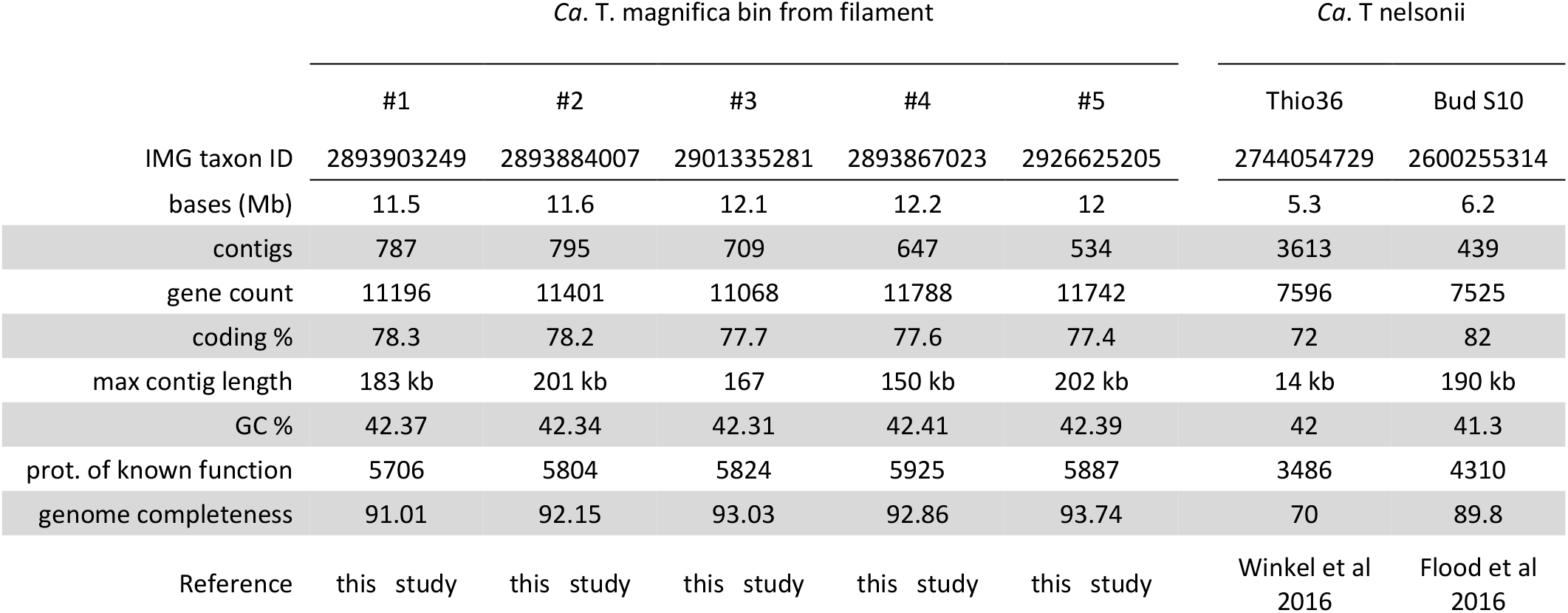
Statistics of the five single cell draft genomes of *Ca*. Thiomargarita magnifica and the two published draft genomes of *Ca*. T. nelsonii.

**Table S5.** Pairwise Average Nucleotide Identity (ANI) between all available *Thiomargarita* genomes. ANI values are colored from green (highest ANIs) to red (lowest ANIs).

|  | Ca. T.<br>magnifica<br>1 | Ca. T.<br>magnifica<br>2 | Ca. T.<br>magnifica<br>3 | Ca. T.<br>magnifica<br>4 | Ca. T.<br>magnifica<br>5 | Ca. T.<br>nelsonii<br>Thio36 | Ca. T.<br>nelsonii<br>Bud<br>S10 |
| --- | --- | --- | --- | --- | --- | --- | --- |
| Ca. T. magnifica 1 |  | 99.63 | 99.84 | 99.85 | 99.84 | 85.23 | 85.33 |
| Ca. T. magnifica 2 | 99.57 |  | 99.68 | 99.68 | 99.63 | 85.30 | 85.35 |
| Ca. T. magnifica 3 | 99.78 | 99.57 |  | 99.87 | 99.83 | 85.17 | 85.32 |
| Ca. T. magnifica 4 | 99.81 | 99.64 | 99.86 |  | 99.86 | 85.22 | 85.33 |
| Ca. T. magnifica 5 | 99.77 | 99.64 | 99.85 | 99.90 |  | 85.39 | 85.46 |
| Ca. T. nelsonii Thio36 | 85.55 | 85.50 | 85.88 | 85.66 | 85.80 |  | 87.07 |
| Ca. T. nelsonii Bud S10 | 85.55 | 85.54 | 85.69 | 85.61 | 85.63 | 86.39 |  |

**Table S6.**
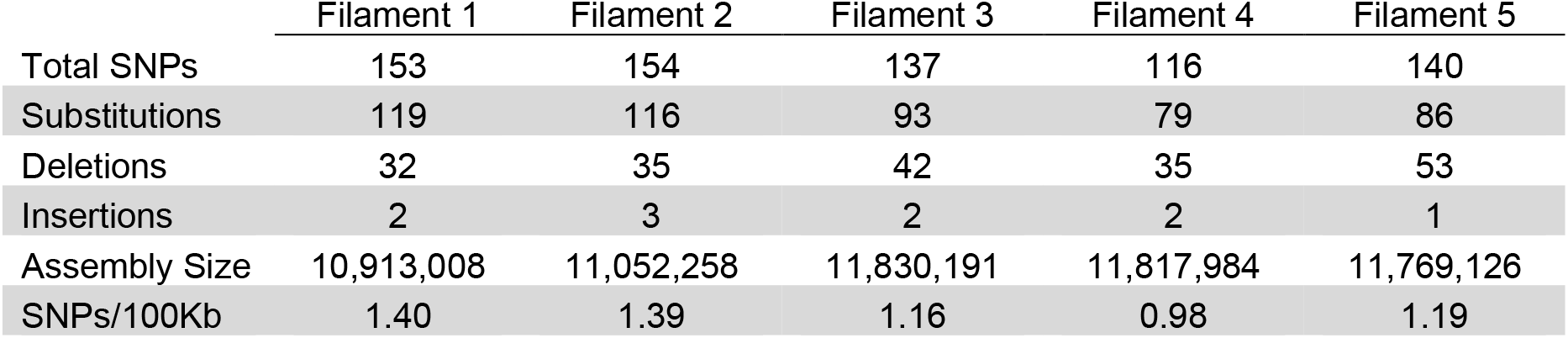
Summary of the variant calling analysis. Detailed information on SNPs in single amplified genomes of five sorted filaments (total number of SNPs and SNPs/100kb) and the breakdown of SNPs by type (substitutions, deletions, insertions).

|  | Filament 1 | Filament 2 | Filament 3 | Filament 4 | Filament 5 |
| --- | --- | --- | --- | --- | --- |
| Total SNPs | 153 | 154 | 137 | 116 | 140 |
| Substitutions | 119 | 116 | 93 | 79 | 86 |
| Deletions | 32 | 35 | 42 | 35 | 53 |
| Insertions | 2 | 3 | 2 | 2 | 1 |
| Assembly Size | 10,913,008 | 11,052,258 | 11,830,191 | 11,817,984 | 11,769,126 |
| SNPs/100Kb | 1.40 | 1.39 | 1.16 | 0.98 | 1.19 |

**Table S7.** Biosynthetic Gene Cluster (BGC) analysis in *Thiomargarita* species, other bacteria (including large sulfur bacteria) and one fungus model system (*Aspergillus nidulans*) famously rich in secondary metabolism.

| genome id | species | large sulfur bacteria | genome size (Mbp) | BGC count | total length of BGCs (Mbp) | percentage of BGCs (%) |
| --- | --- | --- | --- | --- | --- | --- |
| IMG2926625205 | Ca. *Thiomargarita magnifica* fil. 5 | + | 12.01 | 143 | 3.11 | 25.86 |
| IMG2619619276 | *Thioploca ingrica* Lake Okotanpe | + | 4.81 | 50 | 1.01 | 21.04 |
| IMG2600255314 | Ca. *Thiomargarita nelsonii* bud S10 | + | 7.71 | 85 | 1.41 | 18.33 |
| NC_009380.1 | *Salinispora tropica* CNB-440 | - | 5.18 | 16 | 0.90 | 17.30 |
| IMG2772190729 | Ca. *Marithrix* sp. Canyon 246 | + | 3.22 | 30 | 0.54 | 16.70 |
| IMG2788500497 | *Cycloclasticus* sp. SP43 | - | 2.35 | 28 | 0.35 | 15.04 |
| NC_017765.1 | *Streptomyces hygroscopicus* | - | 10.15 | 38 | 1.53 | 15.04 |
| IMG2786546773 | *Thioflexothrix pseukupsi* D-3 | + | 3.97 | 22 | 0.59 | 14.87 |
| IMG2585428147 | *Thiothrix eikelboomii* ATCC 49788 | + | 4.16 | 40 | 0.51 | 12.23 |
| IMG2506520049 | *Thiothrix nivea* JP2, DSM 5205 | + | 4.69 | 45 | 0.57 | 12.13 |
| NZ_CP042324.1 | *Streptomyces coelicolor* A3(2) | - | 8.67 | 27 | 0.98 | 11.35 |
| IMG2515154021 | *Thiolinea disciformis* DSM 14473 | + | 3.92 | 24 | 0.35 | 9.05 |
| GCF_000149205.2 | *Aspergillus nidulans* FGSC A4 | - | 30.28 | 53 | 2.43 | 8.03 |
| IMG2599185268 | *Thiothrix caldifontis* DSM 21228 | + | 3.94 | 24 | 0.30 | 7.71 |
| IMG2619618925 | *Thiothrix* sp. EBPR_Bin_364 | - | 3.75 | 27 | 0.28 | 7.45 |
| IMG2802428810 | *Cocleimonas flava* DSM 24830 | - | 4.49 | 18 | 0.28 | 6.34 |
| IMG2740892603 | *Methylophaga* sp. SM14 | - | 2.89 | 19 | 0.16 | 5.69 |
| IMG2517093012 | *Leucothrix mucor* DSM 2157 | + | 5.19 | 30 | 0.29 | 5.53 |
| IMG2502790011 | *Beggiatoa* sp. Orange Guaymas | + | 4.77 | 26 | 0.26 | 5.38 |
| IMG2556921650 | *Thiothrix lacustris* DSM 21227 | + | 3.72 | 29 | 0.20 | 5.28 |
| IMG2788500402 | *Beggiatoa leptomitiformis* D-401 | + | 4.27 | 18 | 0.18 | 4.17 |
| IMG2515154111 | *Thiofilum flexile* DSM 14609 | + | 3.79 | 23 | 0.16 | 4.15 |
| IMG2508501047 | *Beggiatoa alba* B18LD | + | 4.27 | 16 | 0.15 | 3.60 |
| IMG2684622566 | *Achromatium* sp. WMS3 | + | 3.61 | 10 | 0.11 | 3.17 |
| IMG2236661048 | Ca. *Thiomargarita nelsonii* Thio36 | + | 5.26 | 31 | 0.11 | 2.10 |
| IMG640963011 | *Beggiatoa* sp. PS | + | 6.06 | 13 | 0.07 | 1.12 |

**Table S8.**
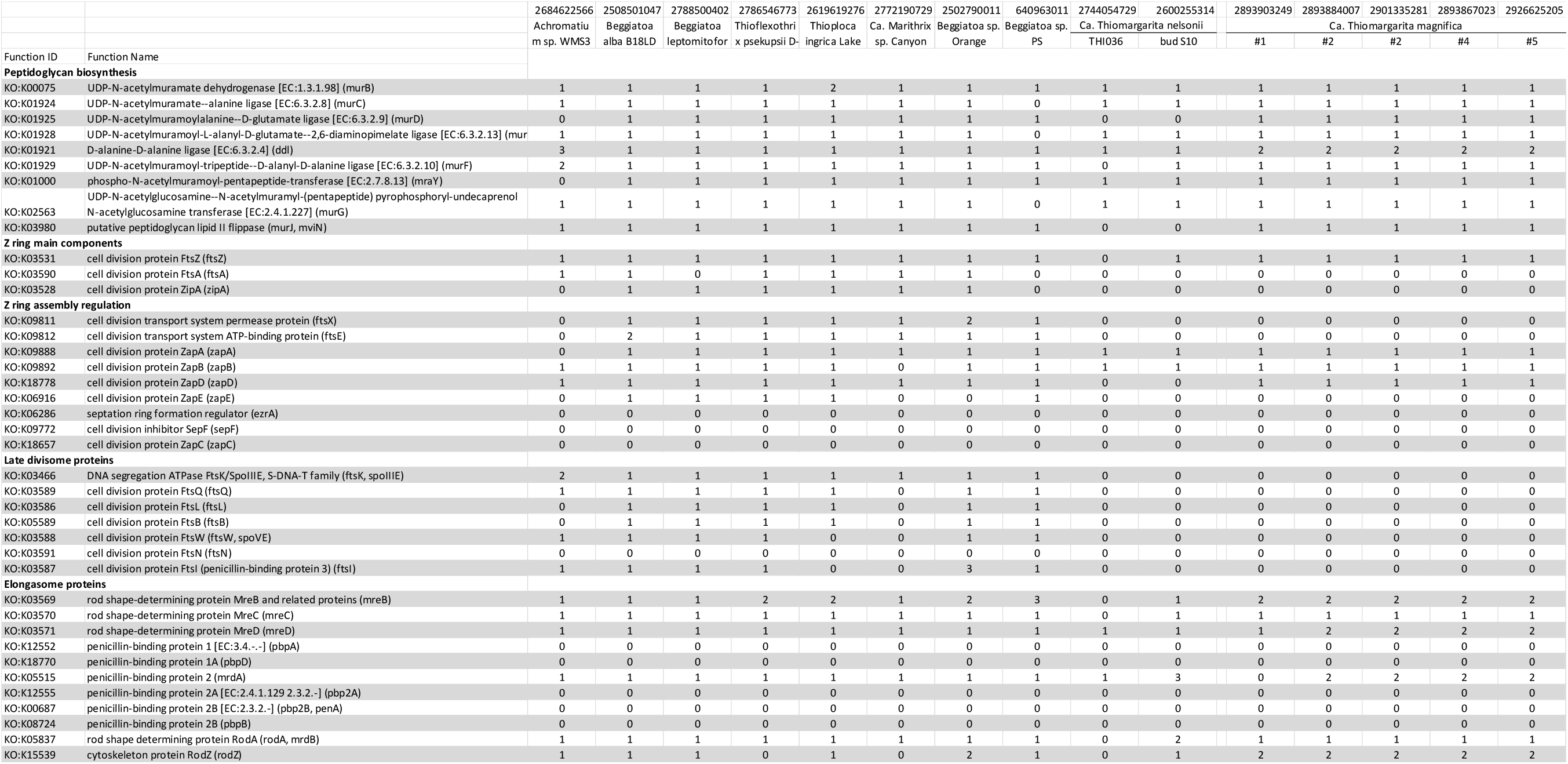
Summary table of the divisome and elongasome genes detected in the genomes of *Ca.* T. magnifica, *Ca.* T. nelsonii and eight other LSBs.

See attached excel sheet for the table below.

**Table S9.** Summary table of the metabolic capabilities of *Ca.* T. magnifica, *Ca.* T. nelsonii and height other LSBs based on their genome analysis.

**Table S10.**
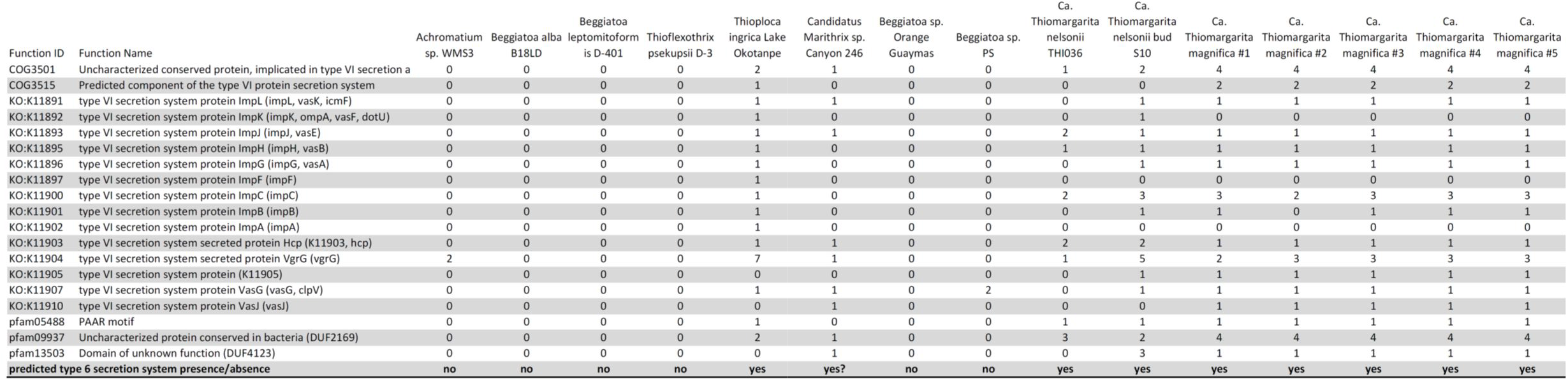
Summary table of type VI secretion genes found in the genomes of *Ca*. T. magnifica, *Ca*. T. nelsonii and eight other LSBs.

**Table S11.**
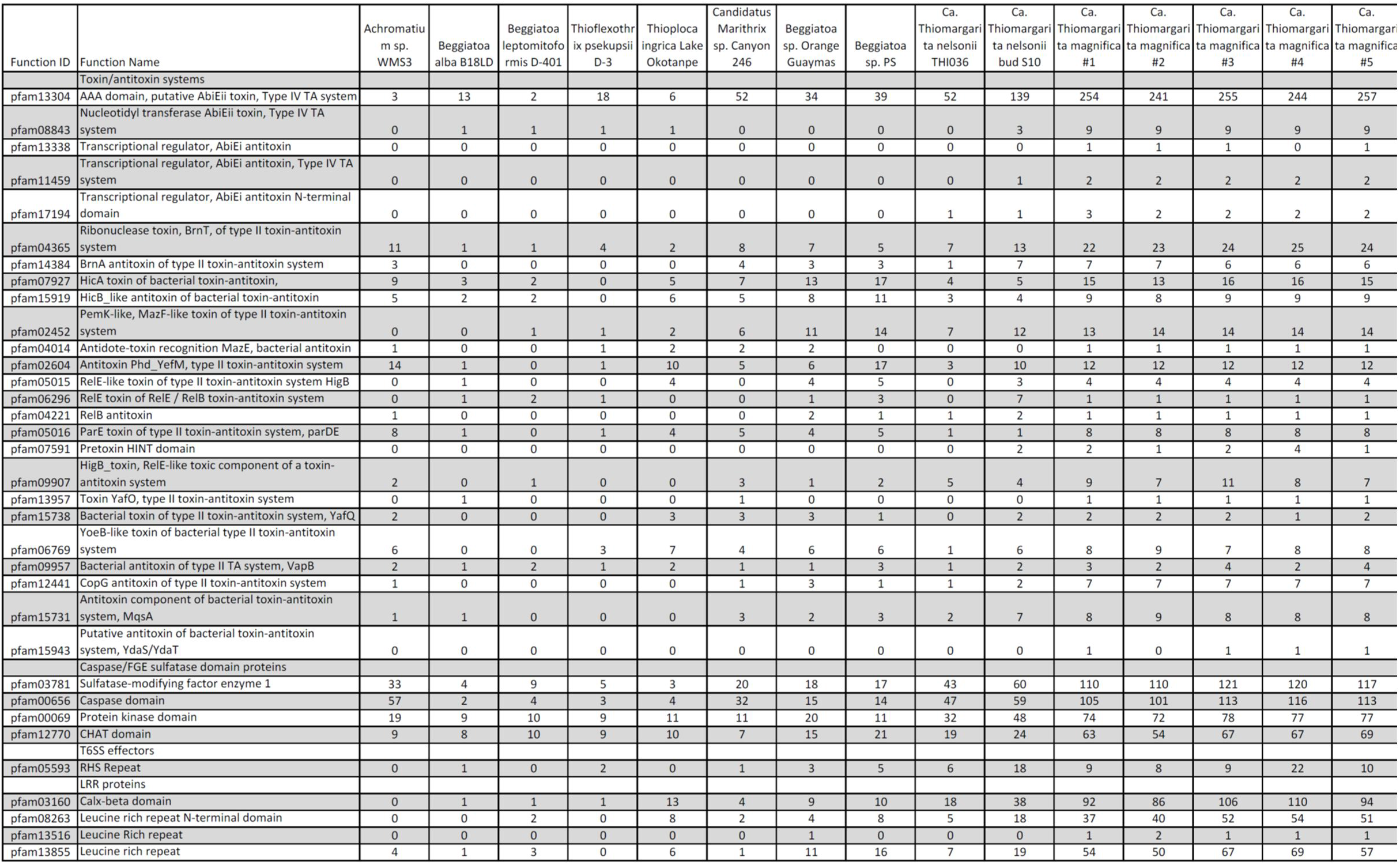
Summary table of Pfam families overrepresented in *Ca*. T. magnifica genomes in comparison to other LSBs.

**Movie S1.**

3D rendering of 6.37 mm long *Ca.* T. magnifica cell observed with hard x-ray tomography (cell *D*, see raw data in Movie S2). The cell cytoplasm has been segmented out and appears continuous along most of the cell length until the most apical constrictions close off completely. See 2D virtual slices in Movie S2.

**Movie S2.**

Fly-through animation of the 6271 virtual slices from cell *D* dataset acquired with hard x-ray tomography with an isotropic resolution of 1 µm. The cell cytoplasm appears as a white ring. Note that samples for HXT were stained with osmium tetroxide to increase contrast on cell membranes (*76, 77*). The cell cytoplasm was segmented on each slice to produce the 3D rendering of the entire cell presented in Movie S1.

**Movie S3.**

Fly-through animation of the virtual slices from the 4.27 mm long cell *G* observed with CLSM after fluorescent labeling of membranes (Table S2). The 3D mesh rendering shows the cell wall in green and the continuous central vacuole in red.

**Movie S4.**

The video shows portions of a *Ca.* T. magnifica and a *Marithrix*-like cell observed in 3D at the confocal laser scanning microscope after staining with the membrane dye FM 1-43x. The membrane septa delimiting the cells of the multicellular *Marithrix-*like filament are clearly visible while no membrane septum is visible inside the *Ca.* T. magnifica cell.

**Movie S5.**

The video shows a portion of the *Ca.* T. magnifica cell used for the estimation of the polyploidy level (cell *M*, Table S2). After DAPI staining, the cell was observed at the confocal laser scanning microscope in 3D by acquiring a z-stack of images. The 3D reconstruction clearly shows the numerous DNA clusters of various sizes spread throughout the cell cytoplasm at the periphery of the central vacuole.

**Movie S6.**

3D rendering of the two smallest cells observed with hard x-ray tomography (cells *A* in yellow and *B* in blue, Table S2). The basal parts of three larger cells (*C*, *E*, and *N*) are also visible. All *Ca.* T. magnifica cells are attached to a sunken leaf and are growing out of the biofilm covering the leaf.

